# Encoding of inflammatory hyperalgesia in mice spinal cord

**DOI:** 10.1101/2021.05.18.444665

**Authors:** Omer Barkai, Rachely Butterman, Prudhvi Raj Rayi, Ben Katz, Shaya Lev, Alexander M Binshtok

## Abstract

Inflammation modifies the input-output properties of peripheral nociceptive neurons, thus leading to hyperalgesia, a condition in which the perception of noxious heat stimuli is altered such that the same stimulus produces enhanced pain. The increased nociceptive output enters the superficial dorsal spinal cord (SDH), which comprises the first CNS network integrating the noxious information. Here we used *in vivo* calcium imaging and a computational approach to investigate how the SDH network in mice encodes the injury-mediated abnormal input from peripheral nociceptive neurons. We show that the application of noxious heat stimuli to the hind paw in naïve mice before induction of injury affects the activity of 70% of recorded neurons by either increasing or suppressing it. Application of the same noxious heat stimuli to hyperalgesic skin following injury leads to activation of previously non-responded cells and de-suppression of the “suppressed” neurons. We further demonstrate that reduction in synaptic inhibition mimics the response to the noxious stimuli in hyperalgesic conditions. Using a computational model of the SDH network, we predict that the “disinhibitory” effect of hyperalgesic stimuli results from the inflammation-mediated increased afferent input to the SDH network and a decrease in SDH inhibition. Both of these processes synergistically contribute to the injury-mediated increase in SDH output towards higher brain centers.

## Introduction

Noxious information is detected by primary afferent nociceptive neurons and transmitted to the superficial laminae of the dorsal spinal cord. There, the information undergoes through a complex neuronal network composed of excitatory and inhibitory interneurons. The output from this network is transmitted via projection neurons to higher brain centers, thus sculpturing the sensation and perception of pain. Tissue inflammation leads to changes in the activity patterns of peripheral nociceptive neurons [4,5,17,70], thus modifying the input and output of the superficial spinal dorsal horn (Laminae I-II, SDH) network [3,10]. While the mechanisms underlying inflammatory-mediated hyperexcitability of nociceptive neurons have been widely studied (see for example [6,7,20,23]), how the SDH network accommodates abnormal input from hyperexcitable nociceptors and how these enhanced inputs affect the SDH network’s output towards higher brain centers, leading to inflammatory pain is not well understood.

Traditionally, noxious stimuli-responsive spinal neurons have been investigated by *ex vivo* preparations or *in vivo* electrophysiology. These studies reveal functional diversity of dorsal horn neurons [9,14,15,24,62,65] and changes in their synaptic connectivity following inflammation or nerve injury [1,60,71]. However, studies describing the response of SDH circuitry to noxious stimuli applied to target organs *in vivo* have been limited. Recently, several studies implemented *in vivo* two-photon laser scanning microscopy (2PLSM) from the spinal cord allowing simultaneous optical recording of the activity of multiple (~200) neurons with a single-cell resolution during noxious stimulation of the skin [25,26,29,55]. This approach permitted for the first time to detail how the SDH network codes thermal stimuli, showing that the heat applied to the skin is represented in a gradual manner, such that application of noxious (above 42°C) stimulus to the skin activates distinct populations of neurons [55]. These results unravel how stimuli are represented by SDH neurons in normal conditions. However, network analysis in pathological conditions, such as hyperalgesia, i.e., enhanced sensitivity to noxious heat stimuli following injury-induced inflammation, is still missing. Here we asked how the stimuli representation in the SDH network changes in inflammatory conditions to produce an altered, hyperalgesic perception of thermal stimuli after injury.

To address this question, we analyzed the changes in the response patterns of SDH neurons to the same noxious stimulus before and after injury *in vivo* in the same animal, using 2PLSM. We show that inflammation promotes an overall increase in neuronal activity of the SDH network. This increase in neuronal activity during noxious heat stimuli under inflamed conditions is predominantly driven by recruitment of silent neurons and a decrease suppression of neurons. Moreover, we demonstrate that pharmacological blockade of SDH inhibitory neurons mimics the effects of injury on the SDH network activity. We implemented our findings into a computational model of the SDH network and demonstrated that a combination of increased afferent input and spinal cord disinhibition could underlie the injury-mediated increased output of the SDH towards higher brain centers. Our results demonstrate how pain-related spinal cord circuitry represents input from the periphery and how this representation changes in pathological conditions underlying the development of inflammatory hyperalgesia.

## Materials and Methods

### Animals

Young adult (4-6 week) C57BL/6J female mice were used for calcium imaging. Usage of young adult females allows relative ease of tissue removed during surgery and minimal attenuation of light intensity during imaging [55]. Mice were group-housed on a 12-h light/dark cycle and were randomly assigned for experiments. All procedures were conducted in accordance with the guidelines and approved by the animal ethics committees of the Hebrew University of Jerusalem (MD18-15608).

### Surgery

The surgical approach was adapted from a method previously reported [25,55]. Accordingly, animals were anesthetized with urethane (2 mg g^−1^), delivered by 2 i.p. injections separated by 30 min. Surgery started 30 min after the second urethane injection. Corneal reflex was examined throughout the experiment, and up to 0.6 mg g^−1^ additional urethane was administered to animals with a corneal reflex response. A tracheotomy was then performed to facilitate smooth breathing and to minimize the drift caused by breathing during the imaging sessions. The core body temperature was maintained at 36°C ± 1°C with a closed-loop heating system throughout the surgery and imaging. The right hind limb was gently shaved with hair removal cream. Paravertebral muscles at vertebrae level T10–L1 were retracted, and spinal clamps (STS-A, Narishige) were used to clamp the exposed vertebral column and stabilize the preparation. A dental drill was used to thin the vertebral column laterally to facilitate the laminectomy. A dorsal laminectomy was then performed using micro scissors (WPI, Sarasota) at vertebra level T12 to expose the spinal cord. A custom-designed plastic chamber was placed around the vertebrae and was sealed with agarose gel (2%) to create a watertight compartment for the use of a water immersion objective. The exposed spinal cord was kept at a stable temperature (27.52 ±0.3°C at the beginning of the experiment and 28.2 ±0.5°C at the end of the experiment, n=3 mice) with normal Ringer solution (in mM: 135 NaCl, 5.4 KCl, 5 HEPES, 1.8 CaCl_2_; pH 7.2). The temperature monitoring was performed using an animal research thermometer (Bioseb). The dura mater was carefully removed with a fine needle, and the animal was rotated around the longitudinal axis by ~30 degrees for imaging. Blood flow through the central vessel was closely monitored as an indicator of tissue health throughout the experiment.

### Dye injections

Neurons in the superficial laminae of the dorsal horn were bulk-loaded with the Oregon Green 488 BAPTA-1 AM (OGB, Invitrogen) under two-photon laser scanning microscopy (2PLSM) as described previously [55]. OGB was chosen over injecting AAV encoding for a genetically encoded calcium indicators to prevent the development of local tissue inflammation, which can occur after local viral injections and might alter the properties of neurons and invading axons [53]. We used glass pipettes with 2–3 μm tips to inject a solution containing 1 mM OGB and 50 μM Alexa Fluor 594 (for pipette visualization [26]), 10% dimethyl sulfoxide and 2% (w/v) Pluronic F-127 (Invitrogen) in Ca^2+^/Mg^2+^ free pipette solution (in mM: 150 NaCl, 2.5 KCl, 10 HEPES; pH 7.4). In some experiments 0.25 μM Sulforhodamine 101 (SR-101) was used instead of Alexa Fluor 594 [44]. The dye solution was then sonicated for 30 min on ice, filtered with a 0.22 μm centrifuge filter (Millipore), and loaded into a glass pipette. The pipette inserted into the picospritzer holder (PV820, WPI, Sarasota, USA) was mounted onto the micromanipulator (SM7, Luigs and Neumann) and controlled via an external module. The pipette tip was targeted to 70–130 μm below the surface of the spinal cord, about 200 μm lateral to the central vessel, using the micromanipulator. Spinal neurons were bulk-loaded for about 3 min using picospritzer (PV820, WPI, Sarasota, USA) by applying 900 ms pulses of 15-25 psi to the pipette to pressure-eject the dye at 3 or more sites approximately 200 μm apart from each other. The dye injection was carried out under a green fluorescent light to check for any blockage in the pipette and to confirm the diffusion of the dye into the tissue using a 10 × dry objective. After dye injection, the imaging site was covered with a No. 0 glass coverslip pre-cut to fit inside the custom chamber, sealed with 2% agarose in Ringer solution, except in the experiments using synaptic blockade pharmacology, in which the imaging site was covered with a plastic coverslip with an access pore for drug application. We waited for 1 h for optimal diffusion of the dye into the cells before commencing the imaging.

### Two-photon laser scanning microscopy imaging

Two-photon calcium imaging experiments were based on Ran et al. [55]. The animal with an exposed spinal cord was placed under a two-photon microscope (Zeiss), and the Ca^2+^ imaging of the dorsal horn was carried out using a Zeiss 20× water-immersion objective (I.R., N.A. = 1.0) with 1× optical zoom. This provided a 438 × 438 μm field of view (FOV) that was scanned at 2 Hz and recorded as a series of 256 × 256 pixels FOV. Alexa Fluor 594, OGB, and SR101 were excited using an 810 nm Ti:sapphire laser (Chameleon, Coherent). Following the optimal dye diffusion, a few reference ROIs in the FOV were randomly picked to check and correct for the spatial drift between the imaging sessions. The SR-101 fluorescence was separated from the OGB fluorescence using a dichroic mirror (562 nm, Semrock Inc., Rochester, NY, USA). Fluorescence emissions were detected simultaneously by two non-descanned photomultiplier tubes with a 542/50 nm filter (Semrock) for ‘green’ fluorescence emission and a 617/73 nm filter (Semrock) for ‘red’ fluorescence emission.

The mouse’s right hind limb paw was placed on a custom-designed 1cm ×1cm Peltier device (**Figure 1A**), with an aluminum tape wrapped around the dorsal side of the paw to keep the paw stretched and stable. The Peltier element was held at a baseline temperature of 32°C before noxious heat stimulus was applied by injecting current with a D.C. power supply (TTi PL-303) digitally tuned and controlled using Digidata 1440A A/D interface (Molecular Devices) and pClamp10 software.

**Figure 1.**
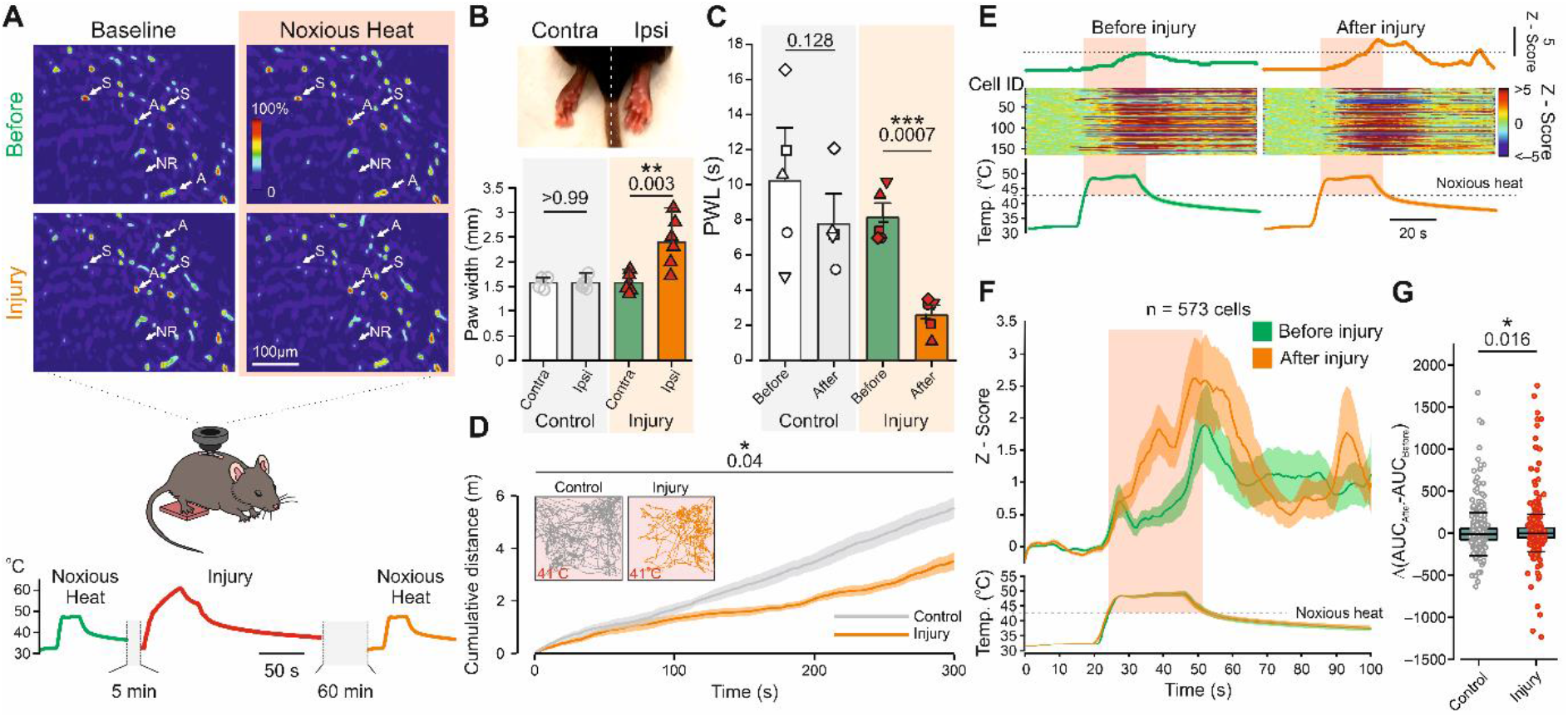
Thermal hyperalgesia is accompanied by increased activation of the superficial spinal dorsal horn (SDH) neurons. **A**. Scheme depicting the experimental procedure. *In vivo* recordings of changes in intracellular Ca^2+^ evoked by noxious heat stimulation (showed in the lower panel) of the right hind paw using a custom Peltier device. *Inset*, pseudocolor photomicrographs of SDH cells before application of the first noxious heat stimulus (*Baseline, Before*), after application of the first noxious heat stimulus (*Before, Noxious Heat*), and before (*Injury, Baseline*) and after application of the second noxious heat stimulus (*Injury, Noxious Heat*) after the burn injury. Arrows outline different response patterns of SDH cells: activated (A), suppressed (S) and non-responsive (NR). **B**. *Upper*, an image of the hind paws contra- and ipsilateral to application of the hyperalgesia-inducing stimulus used in *A* for inducing the burn injury. Note the redness and the edema of the injured hind paw. *Lower*, bar graph and individual values depicting the paw thickness following application of the burn injury to the right hind paw (*Injury*), compared to the contralateral paw and to the control group, which underwent the same procedure but without applying the burn injury. n = 6 mice in each group, unpaired *t*-test. **C**. Bar graphs and paired individual values of paw withdrawal latencies (PWL) measured following focal application of radiant heat before and after application of the burn injury (*Injury*) and in control conditions (*Control*). Paired *t*-test, n=5 mice in the “control” group and n = 6 mice in the “injury” group. **D**. Mean ± SEM of the cumulative distance (measured over 5 minutes) of the exploration of naïve mice (control conditions, *gray*) and mice after application of the burn injury (*orange*) on a warm (41°C) surface. n=6 mice in each group, ordinary two-way ANOVA. *Inset*: representative trajectory traces of naïve (*Control*) and mice following the burn injury (*Injury*). Note a significant decrease in the exploration distance following hyperalgesia-inducing stimulus, suggesting the development of thermal hyperalgesia. **E**. *Middle*, representative activity heat maps for 166 cells in the same field of view (FOV) during application of noxious heat stimulus to the paw before and after induction of hyperalgesia-inducing injury. The averaged changes in activity are shown above the heat maps. The dotted line is aligned to the peak of the mean response before the injury. The stimuli are shown below the heat maps, the noxious (above 42°C) component of the stimulus is outlined by the pink shadow. Representative of 4 mice. For the comparison with the responses in the control conditions, see *Supplementary Figure 1A*. **F**. Changes in intracellular Ca^2+^ following application of the noxious heat stimulus (depicted below) before (*green*) and 1 hour after (*orange*) induction of the burn injury (Z-Score; mean ± SEM) from all recorded cells in 4 mice. The noxious (above 42°C) component of the stimulus is outlined by the pink shadow. n=573 cells from 4 mice. Note an increase in cell activity following the application of the noxious heat stimulus to the injured paw. For the comparison with the responses in the control conditions, see *Supplementary Figure 1B*. **G**. Box plots and individual values of the difference in AUC of responses (AUC_After_ – AUC_Before_) to noxious heat stimulus applied before and 1 hour after induction of the burn injury (*Injury*) and following application of first and second noxious heat stimuli in control conditions (*Control*). The data points were collected and analyzed over the time of the noxious part of the stimulus. Box plots depict the median and 25 and 75 percentiles, and the whiskers depict outlier range. Mann-Whitney test, n_injury_=573 cells from 4 mice; n_control_=630 cells from 4 mice.

For each trial, the spinal cord was imaged for 20 s at the baseline temperature to obtain baseline fluorescence and noise before applying noxious heat stimulus. Two trials of each stimulus were imaged in each spinal FOV. Mild burn injury was induced by delivering a hyperalgesia-inducing (50-65°C) thermal stimulus using the same stimulation source as all other temperature stimuli.

### Image analysis

Videos were imported into ImageJ for concatenation. Movement corrections were performed using a custom-built Matlab code available upon request. Ca^2+^ imaging analysis was performed using the MIN1PIPE calcium imaging signal extraction pipeline [36], which we used for background removal, automatic ROI identification, and signal extraction based on spatial and temporal properties of individual units. A priori structural element estimate was initialized at 5 pixels to seed ROI detection. An average of 145 ROIs were detected in a recording. For population analyses, we calculated Z-scores for the neuronal responses using the mean and the standard deviation of the 20 s baseline period preceding stimulation onsets per each recording session (before vs. after). All traces were smoothed prior to analysis using Matlab’s default Smooth function, with a moving average filter with span = 10. Responses were classified by their ‘before’ traces as “activated” or “suppressed” by the sign of the integral (+/−) during stimulation and if during that period their intensity passed 2 S.D. or −2 S.D. threshold in at least three consecutive time points. Neuronal signals that were not classified as “activated” or “suppressed” and did not pass the threshold value were classified as “NR.” Raw images and individual neuronal responses were visually observed. Experiments or neurons with failed image registration and irregular motion artifacts (because of thermal stimulus-induced paw movement reflex due to insufficient anesthesia) were excluded.

The data points for AUC were collected and analyzed only over the time of the noxious part of the stimulus from 42°C and until 50°C, right after the heat ramp was stopped and the heat started to decrease (“noxious period”). The data were calculated relative to the baseline obtained before the relevant imaging session. No comparison between the baselines of the different recording sessions of the same experiments was performed due to time-dependent sample movement and decay in fluorescence. The area under the curve (AUC) and mean amplitude were measured for each cell.

### Astrocytes activity analysis

The astrocyte activity was detected in separate sets of experiments. To label astrocytes, glass pipettes with 2–3 μm tips containing 1 mM OGB and 0.25 μM Sulforhodamine 101 (SR-101) were targeted 70–130 μm below the surface of the spinal cord, about 200 μm lateral to the central vessel. Astrocytes and neurons were bulk-loaded for about 3 min by applying 900 ms pulses of 15–25 psi to the pipette to pressure-eject the dye at 3 or more sites approximately 200 μm apart from each other. Cells that exhibited both green (OGB-AM) and red (SR-101) fluorescence with a confidence overlap of 80% of the detected cell surface were tagged as astrocytes. Their activity was then analyzed as for neurons (*see above*).

### Behavioral experiments

#### Burn injury induction

Hyperalgesia-inducing heat injury was applied to the glabrous skin surface of the mice’s hind paw with the exact same procedure used in the two-photon experiments by applying 50 to 65°C thermal stimulus for 20s using the Peltier apparatus as described above. The injury protocol was delivered while the animals were deeply anesthetized with isoflurane, to minimize animal suffering and stress. Animals were then returned to their home cages for 1 hour to allow full recovery from anesthesia before returning to the test environment chambers.

#### Warm surface open field test

Burn injury was induced as described above. Animals were placed on a 41°C, sub-noxious heated metal plate surrounded by white plastic walls. Plate size was 20cm x 20cm. The box was cleaned with ethanol and water before each mouse was introduced to the arena. Mice were not introduced to the arena before the experiment. Mice were allowed to explore the arena for 5 min and the behavior was recorded using a video camera for post-hoc analysis. To assess mobility, the distance traveled was analyzed with TrackRodent open-source video tracking software [43].

#### Hargreaves radiant heat test

Two groups (“control” and “injury”) of 6 mice were habituated to handling 48 h before the testing. The mice were habituated to the test environment for 1 h in Plexiglas chambers and the experimental surfaces 24 h before the behavioral tests. The behavioral baseline was obtained by 2 preliminary measurements on the day before and the first day of the experiment. Thermal sensitivity was determined using paw withdrawal latency (PWL) determined with a Hargreaves device (Ugo Basile, Varese, Italy). A radiant heat source was applied to the plantar hind paw at IR=30% intensity. A 20-s cutoff time was used to avoid tissue damage.

#### Tissue thickness measurements

For tissue swelling analysis, a digital micrometer (Mitutoyo) was used to measure the thickness of the plantar area before and at defined time points after injury induction. Thickness increase was calculated as differences from baseline measurements.

#### Computational model

The computational model is composed of 1000 neurons, with 300 inhibitory (Ni, 30%) and 700 excitatory (Ne, 70%) Conductance-based Adaptive Exponential (CAdEx) integrate-and-fire neurons [18,47,48]. 100% of the inhibitory neurons had a tonic firing pattern. Excitatory neurons were divided into three classes based on their firing patterns: delayed (66%), phasic (24%), and single (10%) [52]. Neuronal firing patterns were manually tuned to fit with SDH neuron firing properties according to Prescott et al. [49] (**Table 1**). The aim of the single neuron model was to recapitulate the experimental firing properties of the main classes of SDH neurons. Experimental frequency-current (*f*-I) and delay curves were manually extracted using WebPlotDigitzer.

The equations of a single CAdEx model neuron was as follows:

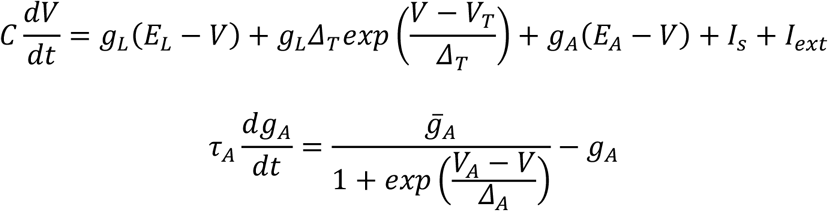

With the following after-spike reset mechanism

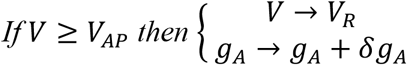

Where *C* represents the membrane capacitance, *E*_*A*_ is the reversal potential of adaptation conductance, *V*_*rest*_ is the membrane’s resting potential, *V*_*A*_ is the subthreshold adaptation threshold, Δ_*A*_ is the slope of subthreshold adaptation. *V*_*T*_ is the spike threshold and Δ_*T*_ is the slope of the spike initiation; *g*_*L*_ is the leak conductance.

An action potential (AP) is initiated when V reaches *V*_*T*_, and the exponential term escalates fast. When it reaches the detection limit (*V*_*AP*_ = 0 *mV*), the membrane potential is set to 30mV and reset to *V*_*R*_, and then goes through a refractory period of *τ*_*ref*_ linearly graduating (slope = 1 · 10^−5^). *g*_*A*_ is incremented by a quantal conductance δ*g*_*A*_ after each spike. *I*_*s*_ and *I*_*ext*_ represent the currents received from synaptic and external (nociceptive) inputs, respectively.

Simulations were computed with time steps of 0.05 ms and lasted 50 s. Model source codes will be available on the ModelDB sharing repository.

### Synapses

The amplitudes of post-synaptic potentials (PSPs) were determined by the spikes emitted by the pre-synaptic neurons in the LIF network. All of the synaptic connections between neurons were unidirectional [37,38] with a connectivity weight, randomly chosen with a 0-to-1 distribution and a mean of 0.5. In every pair of neurons, the unidirectionality (pre- or post-synaptic neuron) was randomly assigned. The input currents’ connectivities for the excitatory populations (E → E, I → E) were enhanced by 2.5-fold [32].

The synaptic connection weights between the neurons are given by the matrix S = (*s*_*ij*_), so that firing of the *j*^th^ neuron instantaneously changes variable *I*_*s*_ of the *i*^*t*h^ neuron 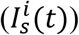 by *s*_*ij*_ :

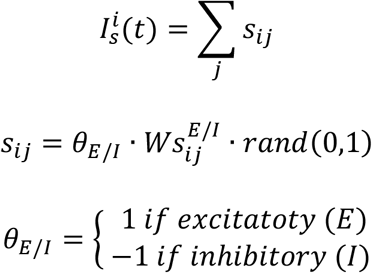

Synaptic weights were tuned to fit neuron-to-neuron pairing experiments [16]) between a pre- (index *i*) and post- (index *j*) synaptic neuron. The excitatory\inhibitory (E\I) weights were 5 · 10^−9^ and 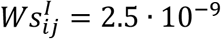; where neuron *i* synaptically projects to neuron *j*, for excitatory and inhibitory neurons, respectively. In the excitatory case, a single AP firing from neuron *i* with a synapse to neuron *j*, lead to an EPSP of ~ 0.5 - 1.5 mv. In the inhibitory case, a single AP leads to an IPSP of ~1-3 mV, values within experimental range values [37]. The sign of the synaptic weight (+/−) was according to E/I characteristic of the neuron, respectively.

### External input

The synaptic weight for the external input to excitatory or inhibitory SDH neurons was *W*_*ext*_ = 50 · 10^−9^ and lead to an EPSP of ~13 mV (or lead to AP firing) agreeing with experimental ranges [28]. To cause a biologically-like variability between synapses, a random variable, ranging between 0 to 1, was introduced to each synaptic connection. The synaptic weights of all external inputs were positive, as nociceptors are excitatory neurons.

Spontaneous activity of SDH neurons has been reported in several *in vivo* [22,30,57] as well as *in vitro* preparations [33,35,55]. To replicate the experimental conditions, we tuned the external input frequencies to published literature EPSC event frequency in SDH neurons. The external inputs were delivered at a designated mean frequency, at random times. All input frequencies stated in the results and the methods section are mean frequencies of random-timed events. While mean external input frequencies received were homogeneous across SDH cells, each cell was excited by an independent and, therefore, different input. A 2 Hz AP input rate for control baseline conditions produced a 7.16 ±0.01 Hz EPSC frequency, and 4 Hz AP input rate for injury baseline conditions produced a 21.93 ±0.04 Hz EPSC frequency, fitting published literature for control vs. inflammatory model conditions [39,40,61,63].

For evoked activity (mimicking heat stimulation in the experimental procedures) in control conditions, the external input frequency was ramped (7 s) to 5 Hz producing a 17.42 ±0.06 Hz EPSC frequency to reach a capsaicin-mediated elevation of EPSC frequency in control conditions in SDH neurons [40]. During the ramp, the mean frequency was gradually increased in 1 s bins. For injury conditions, we increased the external input to 12 Hz, leading to a 61.52 ±0.14 Hz EPSC frequency, reaching the higher rates mentioned in the published literature [2,21]. 70% of randomly chosen SDH neurons received external inputs [64].

For mimicking the disinhibition model, we attenuated the inhibitory synaptic weights to 30% of their original weight values, a value which allowed the maximal inhibition without triggering continuous spontaneous activity reaching the simulations AP frequency saturation values.

### Modeled evoked changes in intracellular Ca^2+^

To mimic calcium imaging integration signals, we used a simplified process of convolution between each neuron’s membrane potential output trace and the following exponential-decay function:

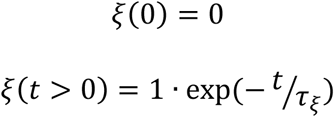

Where *τ*_*ξ*_ = 5 *sec* is the integration time constant, tuned to mimic the calcium imaging traces of the experimental control traces (Figure 1) and then tested on the computational injury and disinhibition models. Each neuron’s intensity integration was then Z – scored with a base line of 7 s. We allowed a 13 s period for the cells to reach a stable baseline. We then classified the neurons to activated, suppressed and non-responsive by comparing the mean intensity integration between baseline and stimulus. When mean intensity during stimulation was higher or lower than baseline, neurons were classified as activated or suppressed, respectively. When equal, they were classified as non-responsive.

### Reagents

GABA and glycine inhibitors, bicuculline methiodide and strychnine (Sigma, St. Louis, MO) were applied by superfusion on top of custom-made plastic coverslips with 3 drug application access pores. Because the coverslip limited the drug diffusion and application time, we used 10 times higher concentrations than previously used in electrophysiological experiments [43,69].

### Statistical analysis

Statistical analyses were performed using Prism 7 (GraphPad). We did not perform a power analysis as the proposed experiments are novel, and cannot estimate the effect size, however sample sizes in the current study are similar to previous reports[29,55]. Mice were randomly allocated to groups in all experiments. Since in some of the experiments comparing AUC data, the data was not normally distributed, non-parametric tests were used for all experiments comparing the AUC data. Mann-Whitney test was used for unpaired values and the Wilcoxon matched-pairs signed-rank test was used for paired values. In the experiments examining the interaction between the nonparametric parameters aligned ranks ANOVA was performed. The data were first aligned-and-ranked using an ARTool (University of Washington) and then ordinary two-way ANOVA was performed on the ranked AUC values. For examining the contingency, either Fisher’s exact test or Chi-square test were used. For the normally distributed behavioral data, unpaired *t*-test, paired *t*-test, ordinary and RM two-way ANOVA were used. Actual p values are presented for each data set. The criterion for statistical significance was p < 0.05. Boxplots presented in all figures depict median, 25^th^; 75^th^ percentile and outlier range. Further information and requests for resources and reagents should be directed to and will be fulfilled by the Lead Contact, Alexander Binshtok.

### Data and code availability

All datasets generated during and/or analyzed during the current study are available in the main text, the supplementary materials, or upon request from the Lead Contact, Alexander Binshtok. Original/source data for Figures 3, 4, 6-8 in the paper will be available on the ModelDB repository.

## Results

### Application of noxious heat stimulus after hyperalgesia-inducing injury leads to increased activity of the SDH network

We examined how superficial spinal dorsal horn (Laminae I-II, SDH) neurons generate a network representation of enhanced input from the peripheral thermosensitive nociceptive neurons, thus encoding inflammation-induced thermal hyperalgesia. We hypothesized that increased input from the periphery would increase the overall responsiveness of SDH neurons. If it is true, would this increase be achieved by enhanced responsiveness of the neurons that responded to noxious heat before the inflammation? Or increased input from the hyperexcitable peripheral neurons would modify the network response signature to noxious heat? To answer these questions, we performed *in vivo* two-photon laser scanning microscopy (2PLSM) imaging to record noxious (~ 49°C) thermal stimuli-evoked calcium signals in a large (~150 neurons/mice) population of laminae I-II SDH neurons, in anesthetized mice, before and after local burn injury (**Figure 1A**). We stimulated the glabrous skin of the right hind paw with a custom-made Peltier device which was held at a baseline temperature of 32°C [47,55,70]. We then applied noxious thermal stimuli by elevating the temperature at a fast rate (~2.5°C/s) to ~49°C and holding it at this temperature for 20 s. During this stimulation, we recorded changes in intracellular Ca^2+^ from Oregon Green 488 Bapta-1 AM (OGB)-labeled laminae I-II SDH cells (**Figure 1A**, *Before*). To examine the changes in SDH neuronal activity following local inflammation, we induced a mild burn injury by applying an intensive thermal stimulus of 50 to 65°C for ~45 s, to the same hind paw using the same stimulation source used for noxious heat application (**Figure 1A**, *below*). A second noxious thermal stimulus of ~49°C was applied, 1 h after generating the injury, to examine changes in SDH network representation of the inflammatory hyperalgesic state (**Figure 1A**, *Injury*). To minimize time-dependent drift in the field of view and photo-bleaching, we allowed only 1 h for the development of inflammation. To assure that the injury protocol and 1-h time window are sufficient to induce injury and inflammatory hyperalgesia, we examined the inflammation signs and pain-related behaviors following the same stimulation paradigm in separate groups of animals. Stimulation of the hind paw with the same intensive, “injury-inducing” stimulus, used in the two-photon apparatus under the same conditions (*see Methods*), produced substantial redness of the glabrous skin and a significant increase in the paw width (**Figure 1B**). Notably, application of the “injury-inducing” stimulus led to a significant decrease in paw withdrawal latency following stimulation with radiant heat, 1 h after injury (**Figure 1C**) and a significant reduction in the active exploration distance at the near-noxious heat threshold (41°C) in the open field test (**Figure 1D**). Altogether, these results suggest that the application of intensive, “injury-inducing” stimulus induces local inflammation accompanied by thermal hyperalgesia, which we hereafter referred to as “hyperalgesia-inducing stimulus.”

Consequently, we utilized the hyperalgesia-inducing stimulus to study the changes in the response of the SDH network to the noxious heat stimulus applied to hyperalgesic skin and compare them to SDH responses in the control conditions, in which the second noxious heat stimulus was applied 1 h after the first noxious heat stimulus but without preceding injury. We show that application of the noxious heat stimulus produced an overall increase in neuronal activity (**Figure 1E, F**, *Before injury*), calculated over the application period of the noxious heat stimuli (*see Methods*). This evoked activity pattern was stereotypical as applying the second noxious heat stimulus 1 h after the first noxious heat stimulus to control animals with no prior tissue injury led to a similar response (p=0.974, Wilcoxon matched-pairs signed-rank test; **Supplementary Figure 1A-C**).

Application of the noxious heat stimulus 1 h after the hyperalgesia-inducing stimulus led to a significant increase in neuronal activity (**Figure 1E, F**, *After injury*, **Figure 1G**).

It is noteworthy that since we are using OGB, the measured activity may not be exclusively attributed to neurons. SDH astrocytes, for example, have been shown to respond to the application of capsaicin [29] and electrical stimulation [11], but not to noxious cold stimuli [55]. To examine the possible contribution of the astrocytes to the changes in the SDH activity, we have performed a separate set of experiments using SR101 together with OGB to label astrocytes. In these experiments, only ~10% of OGB loaded cells were co-labeled with SR101 and activated in response to noxious heat (n=77 astrocytes in 5 mice, on average 15 cells out of 145 cells per experiment). Application of second noxious stimuli following injury did not produce any change in the activity of the astrocytes (**Supplementary Figure 2**). This, together with a relatively low number of labeled astrocytes and considering that some of the SDH neurons are labeled with SR101 [26], suggest that astrocytes contribute minimally to the injury-mediated change of SDH activity.

### SDH neurons respond differently to the noxious heat stimulus

Interestingly, among SDH neurons labeled with OGB, only about 40% of the cells underwent activation following the first noxious heat stimulus, thus we named them “activated.” About 30% of cells did not respond to the first noxious heat stimulus, thus we named them “non-responsive (NR).” Additionally, a notable subset of cells, about 30%, was suppressed by the application of the first noxious heat stimulus, thus we named them “suppressed” (**Figure 1A**, *arrows “A,” “S” and “NR” for activated, suppressed and non-responsive neurons, respectively*). The distribution of the cells to activated, suppressed and NR was similar in the injury and control groups (**Supplementary Figure 3**, p=0.3, between the “control” and “injury” groups, Chi-square test). It is noteworthy that the categorization of cell responses into activated, suppressed, and NR cells does not necessarily overlap with the excitatory and inhibitory nature of SDH neurons as the former describes the cell response, while the latter is its output. Thus, the increase in SDH activity following the input from the inflamed tissue could be achieved by either increasing the responsiveness of the neurons that responded to noxious heat before the inflammation, i.e., “activated” neurons, or by modifying the network response signature to noxious heat by changing the response pattern of the suppressed or NR SDH neurons, or both. To examine whether the inflammation differently affects the responsiveness of activated, suppressed and NR cells, we calculated the difference in the area under the curve (ΔAUC) of the responses of individual activated, suppressed and NR cells before and after the injury and examined the interaction of the cell types and the treatment by comparing it to control conditions. We found that inflammation differentially affects different cell types (p<0.0001, aligned ranks ANOVA, *see Methods*). We next detailed the effect of the inflammation on the various cell types.

### Non-responsive neurons become activated following injury

To examine if the inflammation-mediated increase of the overall SDH network activity results from the recruitment of NR cells, we monitored their activity following the application of the second noxious heat stimulus on the hyperalgesic skin. We show that the second noxious heat stimulus in hyperalgesic conditions leads to a small increase in activity of previously NR cells (**Figure 2A**) which was significantly higher than the change in the control conditions (**Figure 2B**). It is noteworthy that only 48% of the previously NR cells remained NR in inflammatory conditions, 30% became activated while 22% of the cells became suppressed (**Figure 2C**). The change in the activity pattern of the NR neurons under inflammatory conditions was different from the change that occurred in the control conditions: although the percentage of the NR to activated cells was similar, the number of cells that became suppressed was significantly smaller than in the control conditions (22% vs. 48% in control, **Figure 2C**).

**Figure 2.**
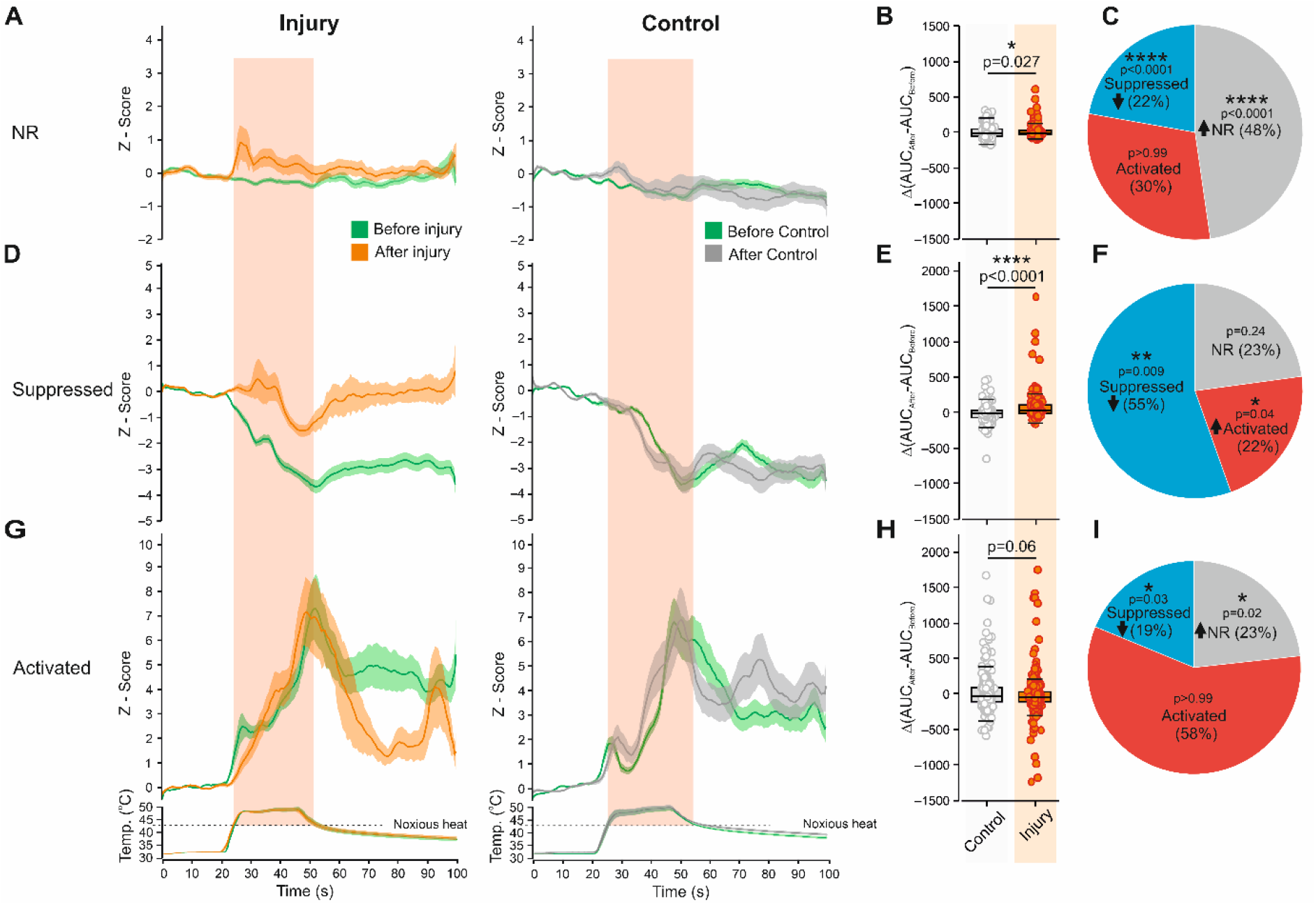
The noxious heat stimuli applied to the hypersensitive skin activates initially non-responsive cells and decrease activity suppression of the initially suppressed cells. **A**. Changes in intracellular Ca^2+^ in all recorded non-responsive (NR) cells from 4 mice, following application of the heat stimulus (depicted below) before and 1 h after induction of the burn injury (*left*) or in control conditions (*right*, Z-Score, mean ± SEM). n_injury_=153 cells from 4 mice; n_control_=185 cells from 4 mice. Note an increase in the activity following the application of the second noxious heat stimulus following the injury, but not in the control conditions. **B**. Box plots and individual values of the difference in AUC of the responses (ΔAUC, AUC_After_ – AUC_Before_) to noxious heat stimulus applied before and 1 h after induction of the burn injury (*Injury*) and in control conditions (*Control*). Mann-Whitney test, n_injury_=153 cells from 4 mice; n_control_=185 cells from 4 mice. **C**. Pie charts depicting the percentage of NR cells which remained NR or became activated or suppressed following injury. The statistical comparison is with the control conditions and the arrow indicates if the number of cells increased or decreased in comparison with the control conditions. Fisher’s exact test. **D**. Same as *A*, but showing changes in the activity of all recorded “suppressed” cells. n_injury_=175 cells from 4 mice; n_control_=188 cells from 4 mice. Note a substantial decrease in the activity suppression following application of the noxious heat stimulus to the injured paw but not in the control conditions. **E**. Same as *B*, but comparing the ΔAUC of the suppressed cells in the injury and the control conditions. Mann-Whitney test, n_injury_=175 cells from 4 mice; n_control_=188 cells from 4 mice. **F**. Same as *C* but depicting the change in the number of the suppressed cells. Fisher’s exact test. **G**. Same as *A*, but showing changes in the activity of all recorded activated cells following the injury and the control conditions. n_injury_=245 cells from 4 mice; n_control_=239 cells from 4 mice. **H**. Same as *B*, but comparing the ΔAUC of the activated cells in the injury and the control conditions. Mann-Whitney test, n_injury_=245 cells from 4 mice; n_control_=239 cells from 4 mice. **I**. Same as *C* but depicting the change in the numbers of the activated cells. Fisher’s exact test. For *A, D*, and *G*, the noxious (above 42°C) component of the stimulus is outlined by the pink shadow. For *B, E* and *H*, the data points were collected and analyzed over the time of the noxious part of stimulus. Box plots depict the median and 25 and 75 percentiles, and the whiskers depict outlier range.

### Hyperalgesic injury promotes disinhibition of the suppressed cells

Another probable explanation for the overall increase of activity in response to noxious heat stimulus applied in the inflammatory conditions might be a change in the activity pattern of the suppressed cells. Indeed, we show that suppressed cells were significantly less suppressed following the second noxious heat stimulus applied to the hyperalgesic skin (**Figure D, E**). We further show that a significantly smaller number of previously suppressed cells remained suppressed following injury (55% vs. 70% in control), and a significantly higher number of suppressed cells become activated (22% vs. 13% in control, **Figure 2F)**.

Collectively, these data imply that the overall increase in SDH circuit activity following hyperexcitable input from the periphery is partly due to the de-suppression of the initially suppressed cells, leading to a decrease in their activity inhibition, and in part by activation of cells that did not respond to the application of the noxious heat stimulus before the injury.

### The computational model of SDH shows the increased activity of the activated neurons in injury-like conditions and suggests that inflammation-induced increased afferent input underlies the altered cell activation following injury

The increase in response to the second noxious heat stimulus, which was applied to the hyperalgesic skin, could also result from an increase in the activity of the activated neurons. Our experimental model, however, does not show any change in the overall activity of the activated neurons following injury (**Figure 2G, H**). Interestingly, we show that although the percentage of the activated cells that remained activated does not change following injury (58% vs. 57% in control), a significantly smaller number of activated cells become suppressed (19% vs. 27% in control) such that a significantly higher number of activated cells become NR (23% vs. 15% in control, **Figure 2I**). The decreased number of activated-to-suppressed cells after injury compared to control implies that following injury, the overall activity of the activated cells might increase. We, however, did not detect any difference between the injury and the control groups (**Figure 2H**). These results are surprising considering the previous studies (*see, for example* [58]). The lack of the change in the activity of the activated neurons may result from the relatively low acquisition rate (2 Hz) we used to sample the data due to a technical limitation of our 2PLSM system. Since a low acquisition rate does not allow recording in high temporal resolution, the resulting calcium signals only allow interpretation of changes relative to a global baseline, before vs. after stimulation, and does not identify changes in spontaneous baseline activity, which may also be affected by the inflammation-induced nociceptive hyperexcitability [7,70]. Moreover, due to photo-bleaching, we were unable to perform continuous recording during the whole experiment, which takes more than 2 h. Instead, we recorded only a short baseline followed by a stimulus-induced response in both conditions. Therefore, we were unable to normalize the response to the noxious heat stimuli in the inflammatory conditions to the baseline at the beginning of the experiment due to the time-dependent changes in cell position and fluorescence. Instead, we calculated the amplitude of the response in the inflammatory conditions by normalizing it to the baseline we obtained in these conditions. It is plausible that this baseline is higher as a result of inflammation-mediated increase bombardment of the SDH neurons from the peripheral neurons. Because we measured the increase in the response amplitude relative to the global baseline, the resulting normalized response to the higher baseline may mask the real magnitude of the observed effect, thus underestimating the actual changes in the activity of SDH neurons, primarily of the “activated” neurons.

To examine the changes in the activity of SDH neurons, taking into account the increased baseline, we modeled the SDH network utilizing a Conductance-based adaptive Exponential (CadEx) [18] integrate-and-fire model (**Figure 3A**). This model allows performing large network simulations with simplified models reproducing neuronal firing properties [18]. We constructed a biologically constrained model of a superficial spinal dorsal horn laminae I-II, composed of 70% excitatory and 30% inhibitory spiking neurons [48,51,62] connected to each other (**Figure 3A, B**). The intrinsic connectivity of the excitatory and inhibitory neurons within the network was unidirectional [37,38], and the synaptic weights of excitatory and inhibitory synapses connected to excitatory neurons were higher than that connected to inhibitory neurons [32] (*see Methods*).

**Figure 3.**
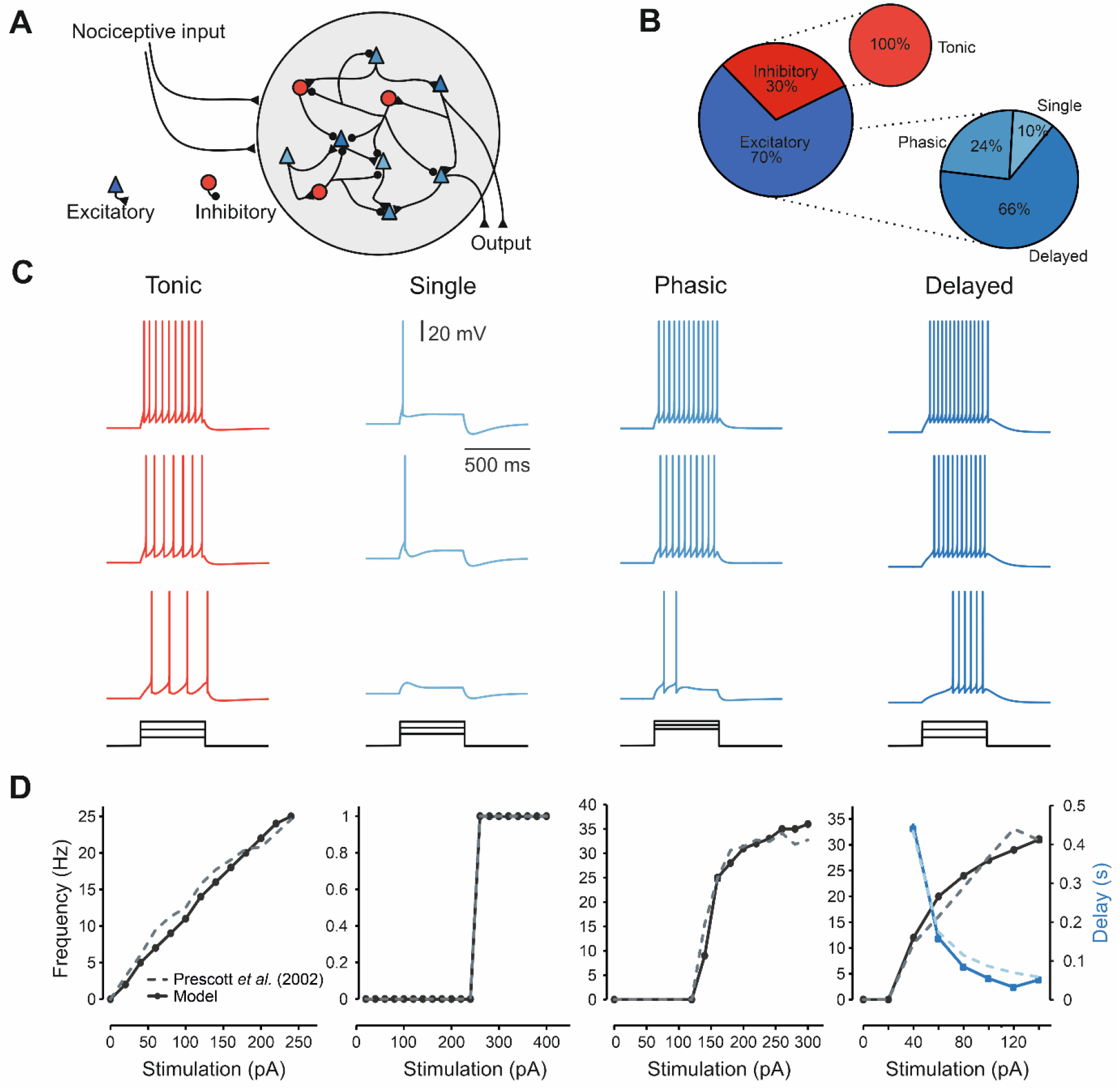
SDH network model. **A**. Scheme depicting the modeled network, which consists of 1000 neurons including inhibitory (30%, red circles) and excitatory local interneurons and projections neurons (70%, blue triangles), interconnected to each other. The network receives nociceptive input simulated by EPSC-like currents at various frequencies, depending on the modeled conditions. In simulated control conditions, 2 Hz input models the baseline activity and 5 Hz input models the activation of nociceptive neurons by noxious heat. The injury conditions are simulated by currents at 4 Hz for baseline and at 12 Hz for evoked activation. **B**. Pie charts showing the percentage of the inhibitory and excitatory cells composing the network and their distribution according to the firing pattern. **C**. Modeled firing of tonic, single, phasic, and delayed firing-type SDH neurons under escalating current clamp steps. Shades of blue represent the modeled excitatory neurons, red represents modeled inhibitory neurons. **D**. Modeled frequency-intensity (*f-*I) curves for each type of neuron (*continuous line*) compared with its respective experimental *f-*I curve from Prescott and De Koninck (2002, *dashed line*). In the delayed neuron, the delay (time to first AP) is also compared with the experimental data.

To capture basic spiking features of SDH dynamics, we modeled three well-established types of excitatory neurons with phasic, delayed, and single firing pattern properties and one type of tonically firing inhibitory neuron [52], **Figure 3C**). The modeled firing properties of these neurons closely resembled the experimental frequency-current (*f-I*) curves and the ‘delay curve’ of the delayed neurons [50] (**Figure 3D**). We simulated the SDH circuitry using a network of 1000 neurons, of them 300 inhibitory neurons, of which all had tonic firing patterns, and 700 excitatory neurons, of which 25% had phasic, 12.5% delayed, and 62.5% single firing patterns [52] (**Figure 3B**). In addition to the intrinsic excitatory and inhibitory connections, excitatory and inhibitory neurons also received excitatory inputs with controlled frequency but at random times, simulating nociceptive input from the periphery during baseline and heat stimulation (*see Methods*). To simulate naïve conditions, we stimulated the SDH network model with input currents at 2 Hz and 5 Hz to model baseline and “noxious heat” induced circuit activity, respectively. These stimulation frequencies generate simulated EPSC frequencies in the SDH model, which were similar to the baseline and evoked SDH EPSC frequencies reported *in vivo* [39,40,61,63] (*see Methods*), thus mimicking the SDH circuit activity before and during noxious stimulation, respectively. Tissue injury leads to an increase in both the spontaneous and evoked activity of the peripheral nociceptive neurons [7,70]. Thus, to reflect the increased spontaneous activity of the hyperexcitable primary afferents, to simulate injury-like conditions, we increased the frequency at the baseline conditions to 4 Hz [40], *see Methods*). To simulate the increased evoked activity during the heat stimulus, we increased the input currents frequency to 12 Hz [2,21] (*see Methods*).

We utilized a calcium imaging-like module, which as in calcium imaging, predicts integration dynamics of the electrical neuronal network to examine whether the simulations qualitatively emulate our experimental data (*see Methods*). Stimulation of the model at the simulated naïve conditions led to an overall increase in neuronal activity (**Figure 4A**, *left, solid green*). Analysis of the individual modeled cells showed that similar to what we demonstrated *in vivo*, the majority of the cells (44%) increased their activity during the modeled stimulus (activated), about 20% decreased their activity (suppressed), and the rest did not change their activity (NR) during the application of the simulated noxious heat stimulus (**Figure 4A**, *inset*). Analysis of the activity patterns of the different neuronal types showed that stimulation with the simulated noxious heat stimulus in control condition activated the majority of the inhibitory tonic (73%) and excitatory phasic simulated neurons (65%), whereas most of delayed and single firing excitatory neurons were non-responsive (62% and 97% respectively, **Figure 4B**). Since our model accurately replicates the activity of the SDH network measured *in vivo*, we have decided to utilize this model to examine the possible mechanisms of the injury-mediated changes in the SDH activity.

**Figure 4.**
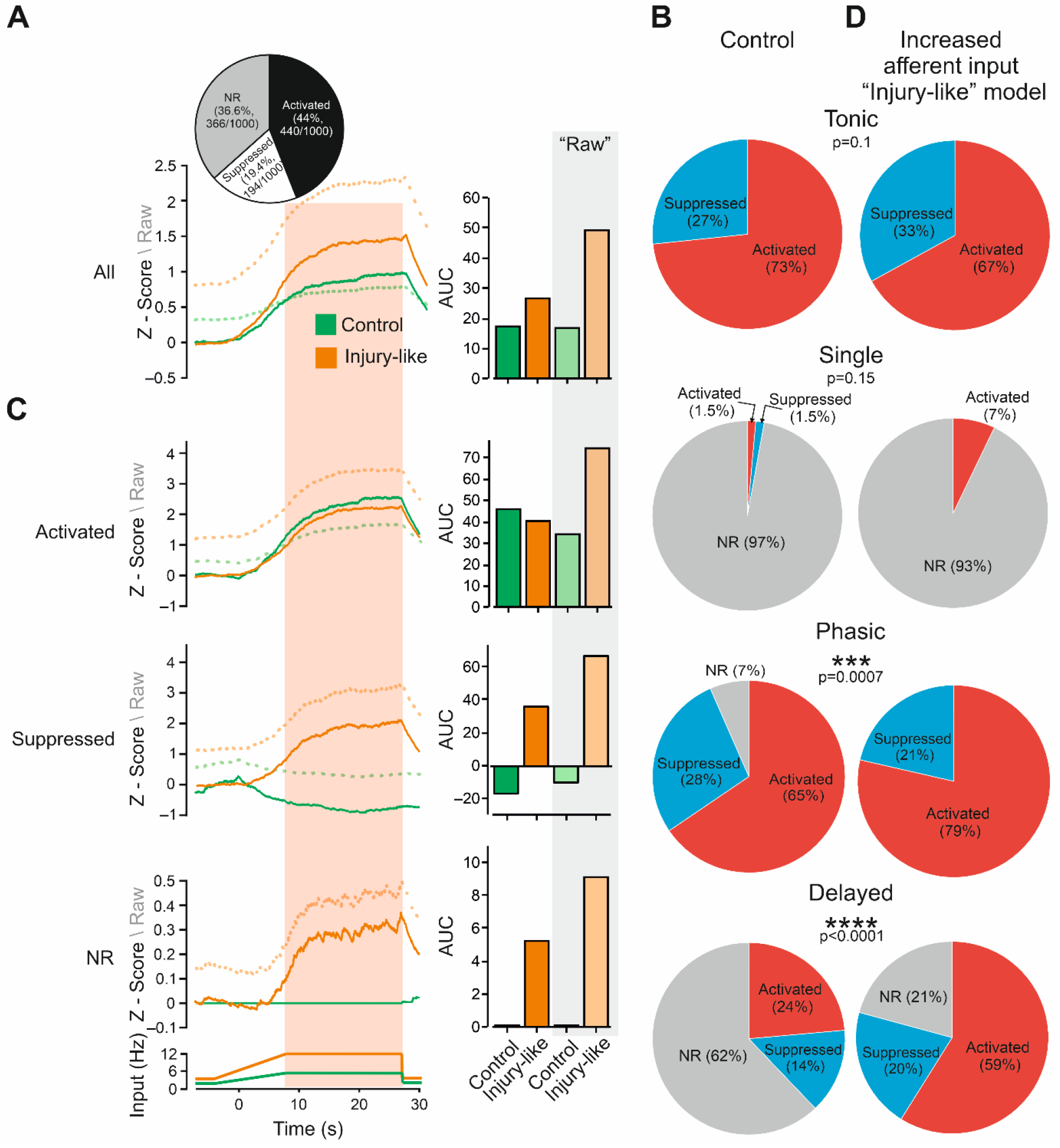
SDH network model predicts an increase in activation of activated and NR neurons and de-suppression of suppressed neurons following increased afferent input. **A**. *left*, mean neural activity integration (1000 modeled neurons) in response to a simulated noxious heat stimulus applied in control (*green*) and increased afferent input (*injury-like, orange*) modeled conditions. The solid lines depict the Z-scored changes; the dotted lines depict the unnormalized “raw” changes in the activity integration between the modeled control and injury-like conditions. *Inset*. The pie chart depicts the fraction of activated, suppressed, or NR neurons. *Right*, Bar graphs of AUC of responses (shown in *left*) to the simulated noxious heat in modeled control and increased afferent input (injury-like) conditions calculated using Z-score (*green* and *orange bars* for control and injury-like conditions, respectively) and “raw” (unnormalized) changes (*light green* and *light orange bars* for control and injury-like conditions, respectively). **B**. Pie charts depicting the proportion of activated suppressed and NR neurons in modeled tonic, singe, phasic, and delayed firing neurons following simulated noxious stimulation in control conditions. **C**. same as *A*, but for activated, suppressed and NR neurons. Note y-axis expansion for simulated NR neurons. **D**. Same as *B*, but in simulated inflammatory conditions. The *p* values are for the comparison with the control (shown in *B*), Fisher’s exact test for tonic firing neurons and Chi-square test for single, phasic, and delayed firing neurons, comparing to the control conditions. n_Tonic_=300 neurons, n_Single_=70 neurons, n_Phasic_=168 neurons, n_Delayed_=462 neurons.

We showed that stimulation of the model by the increased input to simulate the increased evoked activity during the heat stimulus, without affecting the synaptic connectivity between the SDH neurons, leading to an increase in overall activity (**Figure 4A**). These results suggest that the increase in the afferent input is sufficient by itself to enhance the activity of the SDH network.

Similar to the *in vivo* results, the increase in modeled SDH following simulated enhanced afferent input resulted from the activation of the suppressed and NR neurons, but not activation of the activated neurons (**Figure 4C**, *left, solid lines*, **Figure 4C**, *right*). However, by analyzing the activity of simulated SDH neurons, taking into account simulated inflammation-induced changes in the baseline activity, we show substantial activation of the activated cells together with activation of the suppressed and NR neurons (**Figure 4C**, *left, dotted lines*, **Figure 4C**, *right*). Importantly, taking into consideration inflammation-induced changes in the baseline activity, we show that the overall increase in activity becomes substantially larger (**Figure 4A**, *left, dotted lines*, **Figure 4A**, *right*), implying that Z-scoring which we performed for the *in vivo* experiments may mask the extension of the activation of the activated neurons and thus may underestimate the increase in overall activity following inflammation. These results suggest that increased afferent input leads to enhanced activation of activated neurons.

Further analysis of the increased afferent input-mediated changes in the activity patterns of different neurons shows that application of the simulated noxious stimulus in the simulated injury model does not affect the number of activated or suppressed inhibitory tonic neurons but leads to a significant increase in the number of activated excitatory phasic and delayed neurons at the expense of the NR phasic and delayed neurons (**Figure 4D**).

These results suggest that an increase in afferent input replicates well the injury-mediated changes in SDH activity. It also predicts that an increase in spontaneous (background) activity contributes to the overall increase in SDH activity by enhancing the activity of activated neurons.

Altogether our experimental and computational results suggest that SDH circuitry encodes enhanced input from the injured tissue in a complex combinatorial manner, i.e., by changing the activity patterns of the engaged neurons and affecting previously non-responding neurons.

### Blockade of the spinal cord inhibition mimics the effect of an injury

Our results show that the input from the periphery during noxious heat stimuli under naïve conditions produces a reduction in the activity of neurons (suppressed neurons). These results suggest the existence of an inhibitory drive during noxious heat stimuli in the SDH circuity. Furthermore, we showed that in hyperalgesic conditions, the noxious heat stimuli lead to a decrease in the activity suppression of the suppressed cells and activation of a subset of previously NR cells, thus inducing the overall activation of the SDH circuitry, suggesting that the inhibitory drive under this condition is reduced. To test whether inflammation-induced disinhibition of SDH may be an additional mechanism for the injury-mediated increase in SDH activity, we examined the response of SDH cells to noxious heat stimuli following the blockade of inhibitory synaptic inputs at the SDH. To this end, we tested the response to noxious heat stimuli before and after local application of bicuculine (BMI, 200 μM) and strychnine (40 μM) to block GABA and glycine receptors, respectively. These concentrations were used at in order of magnitude above the previously used concentrations to assure that the pharmacological reagents would reach the neurons through the plastic imaging cover window [43,69].

We show that the application of BMI and strychnine leads to an overall increase in the activity of the SDH neurons (**Figure 5A-C**). To explore the mechanism of this increase in activity, we examined the effect of disinhibition on the activity of activated, suppressed, and NR cells. The distribution of cells to activated, suppressed, and NR before the application of BMI and strychnine was similar to the control conditions shown in Supplementary Figure 3 (**Figure 5D**, p=0.25, Chi-square test). Importantly, the comparison of ΔAUC of individual activated, suppressed, and NR cells treated with BMI and strychnine to ΔAUC of activated, suppressed, and NR cells in the control group shows that the effect of disinhibition on the overall activity results from the differential effect of the disinhibition on the different cell types (p<0.0001, aligned ranks ANOVA, *see Methods*). As expected, treatment with BMI and strychnine reversed the activity suppression of the suppressed cells (**Figure 5E, F)**, such that a significantly smaller number of previously suppressed cells remained suppressed following injury (32% vs. 70% in control), becoming either NR (31% vs. 17% in control) or activated (37% vs. 13% in control, **Figure 5G**). Pharmacological disinhibition also led to an overall activation of the NR cells (**Figure 5H, I**) likely by significantly decreasing the number of NR cells which became suppressed (19% vs. 48% in control). Since the overall number of NR-to-activated transitions remained unchanged, the pharmacological disinhibition likely increased the transition from the suppressed to the NR cells (**Figure 5J**).

**Figure 5.**
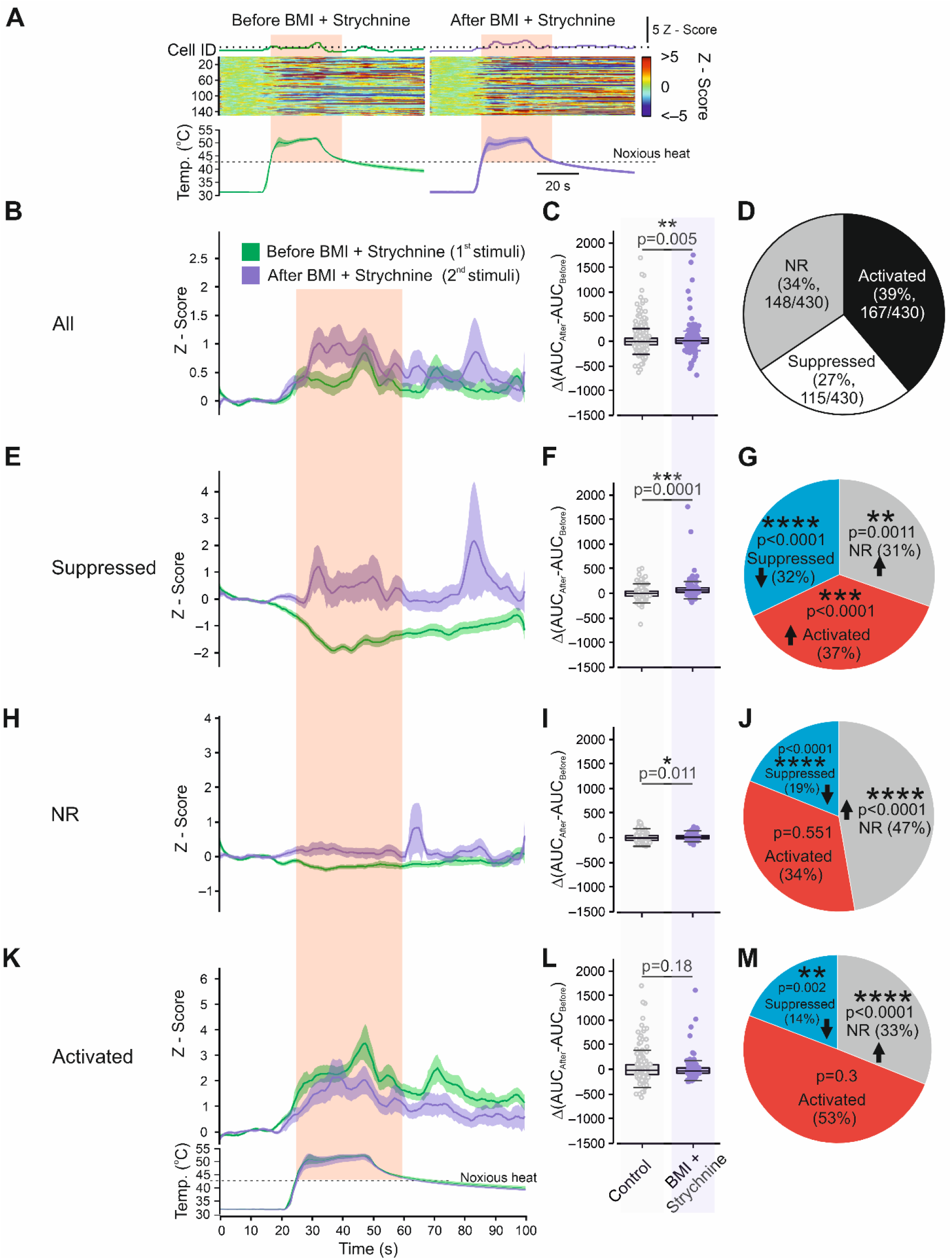
Blockade of the SDH inhibitory synapses mimics the injury-mediated hyperalgesic response of the SDH neurons. **A**. Representative heat maps of the activity of all recorded cells in the same FOV during application of noxious heat stimulus to the paw before and 1 h after treatment with 200 µM bicuculine (BMI) and 40 µM strychnine, to block GABA and glycine receptors, respectively. The stimuli are shown below the heat maps. The averaged changes in activity are shown above the heat maps. The dotted line is aligned to the peak of the mean response before the drug application. n=149 cells. Representative of 3 mice. **B**. Changes in the intracellular Ca^2+^ (Z-Score, mean ± SEM) of all recorded cells before (*green*) and after (*magenta*) application of BMI and strychnine. Note a substantial increase in the overall activity of SDH cells. n=430 cells from 3 mice. **C**. Box plots and individual values of the difference in AUC of responses (ΔAUC, AUC_After_ – AUC_Before_) to noxious heat stimulus applied before and 1 h after application of bicuculine and strychnine (*BMI + strychnine*) and following application of first and second noxious heat stimuli in control conditions (*Control*). Mann-Whitney test, n_BMI + Strychnine_=430 cells from 3 mice; n_control_=630 cells from 4 mice. **D**. Pie chart depicting the fractions of NR, activated and suppressed cells in response to the first noxious heat stimuli in the “BMI + strychnine” group before application of the drugs. **E**, Same as *B*, but showing changes in the activity of all recorded “suppressed” cells. n_BMI + Strychnine_=167 cells from 3 mice; n_control_=188 cells from 4 mice. Note a substantial decrease in the activity suppression following application of BMI and strychnine. **F**. Same as *C*, but comparing the ΔAUC of the suppressed cells in the “BMI + strychnine” and the control conditions. Mann-Whitney test, n_BMI + Strychnine_ =167 cells from 3 mice; n_control_=188 cells from 4 mice. **G**. Pie charts depicting the percentage of suppressed cells which remained suppressed or became activated or NR following application of BMI and strychnine. The statistical comparison is with the control conditions and the arrows indicate if the number of cells increased or decreased in comparison with control conditions. Fisher’s exact test. **H**. Same as *B*, but showing changes in the activity of all recorded “NR” cells. n_BMI + Strychnine_=148 cells from 4 mice; n_control_=185 cells from 4 mice. **I**. Same as *C*, but comparing the ΔAUC of the NR cells. Mann-Whitney test, n n_BMI + Strychnine_=148 cells from 3 mice; n_control_=185 cells from 4 mice. **J**. Same as *G*, but depicting the change in the numbers of the NR cells. Fisher’s exact test. **K**. Same as *B*, but showing changes in the activity of all recorded “activated” cells. n_BMI + Strychnine_=115 cells from 3 mice; n_control_=239 cells from 4 mice. **L**. Same as *C*, but comparing the ΔAUC of the activated cells. Mann-Whitney test, n_BMI + Strychnine_=115 cells from 3 mice; n_control_=239 cells from 4 mice. **M**. Same as *G* but depicting the change in the numbers of the activated cells. Fisher’s exact test. For *B, E, H* and *K*, the noxious (above 42°C) component of the stimulus, is outlined by the pink shadow. For *C, F, I* and *L*, the data points were collected and analyzed over the time of the noxious part of stimulus. Box plots depict the median and 25 and 75 percentiles, and the whiskers depict outlier range.

Our experimental results did not show a change in the activity of the activated cells following pharmacological disinhibition **(Figure 5K, L**). However, we did show that the number of activated cells which become suppressed is significantly smaller (14% vs. 27% in control, **Figure 5M**), suggesting a possible increase in activity of the activated cells, which is masked by the Z-scoring, as described above. To examine the effect of disinhibition on the activated neurons considering the change in baseline activity, we simulated a decrease of the inhibitory inputs (70%) applied to the SDH network model (**Figure 6**). Similar to the *in vivo* results, our model showed that the decrease in the SDH inhibition leads to an increase in the overall evoked integrated activity compared to control AUC (17.1 ± 1.2 vs. 30.3 ± 2.1, **Figure 6A**) by activation of the NR neurons (AUC_Control_ = 0 vs. AUC_Disinhibition_ = 3.4 ± 0.2) and a decrease in the activity suppression of the suppressed cells (AUC_Control_ = –16.6 ± 0.7 vs. AUC_Disinhibition_ = – 5.3 ± 0.7, **Figure 6B**). Importantly, our model predicts that disinhibition leads to a substantial increase in the activation of the activated cells (AUC_Control_ = 34.3 ± 1 vs. AUC_Disinhibition_ = 57.8 ± 1.5, **Figure 6B**). The analysis of the changes in the activity patterns of inhibitory tonic and excitatory single firing, phasic, and delayed neurons shows that, similar to the injury-like model, the application of simulated noxious like stimuli under reduction of SDH inhibition does not lead to changes in the number of activated or suppressed tonic neurons (**Figure 6C**). Unlike the injury-like model, the disinhibition was insufficient to increase the number of activated phasic excitatory neurons, nevertheless, it significantly increased the number of activated delayed neurons at the expense of the delayed NR neurons (**Figure 6C**). Altogether these results suggest that changes in the activation pattern of the pain-related SDH network, which encodes thermal hyperalgesia, may result at least in part from the acute decrease in the inhibitory drive.

**Figure 6.**
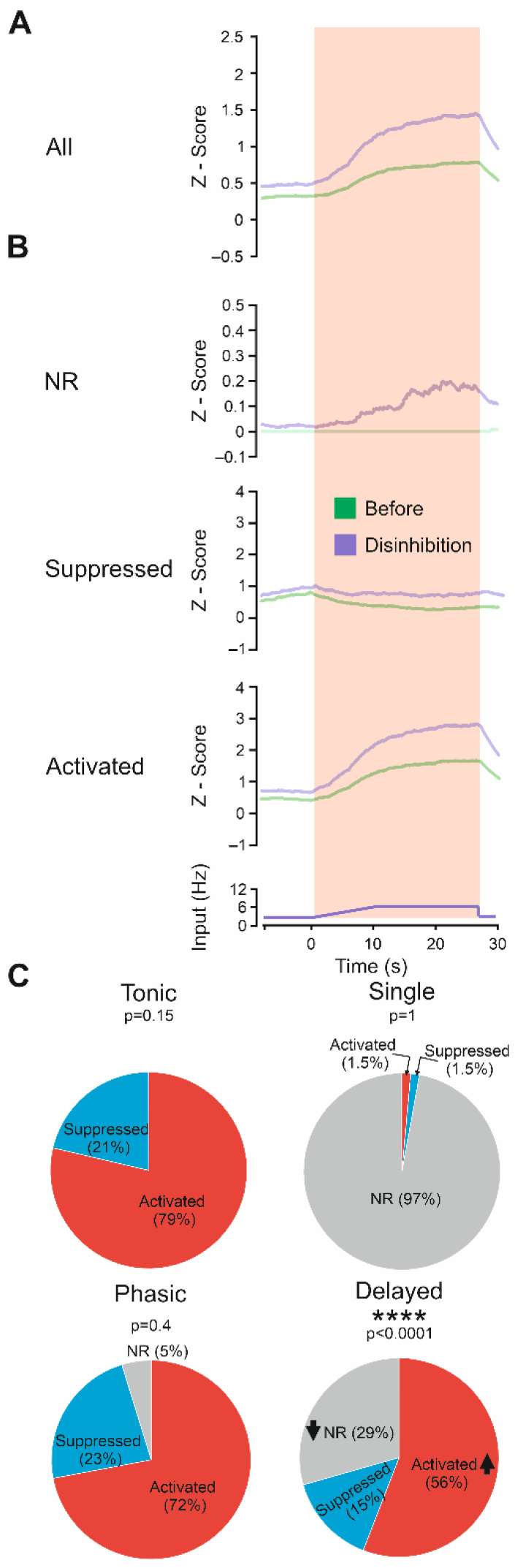
SDH network model predicts an increase in activation of activated and NR neurons following a decrease in SDH inhibition. **A**. Mean of neural activity integration (1000 modeled neurons, “raw” unnormalized changes), in control (*green*) and disinhibitory (*magenta*) modeled conditions. **B**. Mean activity (“raw” unnormalized changes) of the simulated NR, suppressed and activated neurons in simulated control (*green*) and decreased in inhibition (70%, *magenta*) conditions. Note y-axis expansion for simulated NR neurons. **C**. Pie charts depicting the proportion of activated suppressed and NR neurons in modeled tonic, singe, phasic, and delayed firing neurons in response to simulated noxious stimulation following modeled disinhibition. Fisher’s exact test for tonic neurons and Chi-square test for single, phasic, and delayed firing neurons, comparing to the control conditions (*shown in Figure 4B*). n_Tonic_=300 neurons, n_Single_=70 neurons, n_Phasic_=168 neurons, n_Delayed_=462 neurons.

### The computational SDH model predicts that injury increases the activity of excitatory SDH neurons

Our *in vivo* results demonstrated that “noxious heat” stimuli in naïve and inflamed conditions increase the overall neuronal activity. However, it is unclear whether this is translated into an increase in the excitatory output of the SDH network. Our data imply that the enhancement of overall neuronal activity may involve increased activity of both excitatory and inhibitory neurons, which may have unpredictable net effects. We, therefore, used the SDH network model to predict how changes in the input affect the activity of individual excitatory and inhibitory neurons (**Figure 7A, B**), focusing on the population of excitatory neurons with either phasic or delayed firing patterns as they are strongly associated with the projection neurons [19,34,56].

**Figure 7.**
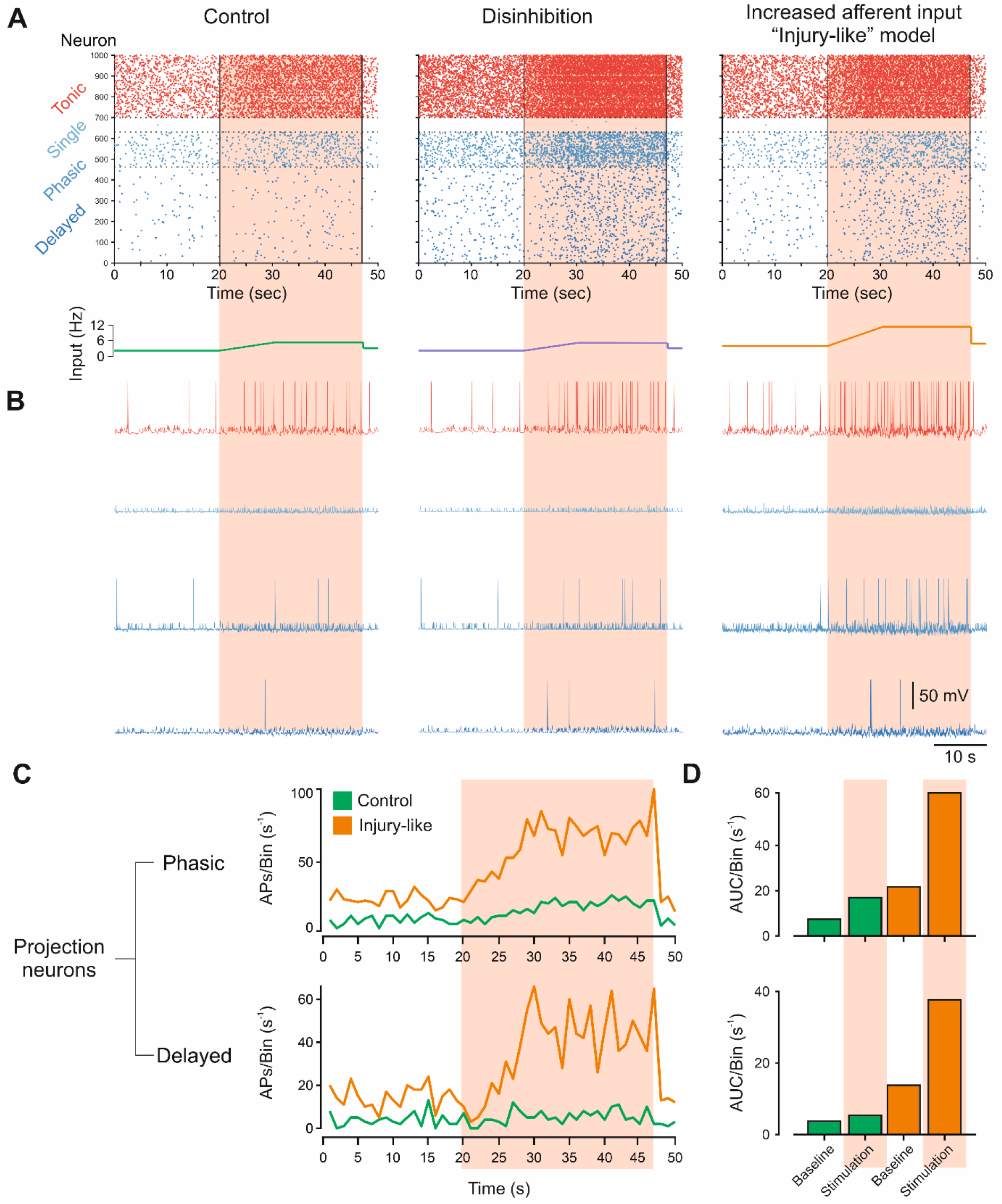
Modeled inflammatory conditions predict an increase in the firing of excitatory SDH neurons. **A**. Raster plots of 1000 spiking neurons in modeled control, disinhibition, and injury-like (increased afferent input) states. Neurons are grouped by their intrinsic firing patterns and colored according to their excitatory (*delayed, phasic, single in shades of blue*) or inhibitory (*tonic in red*) properties, respectively. The stimulation frequency is depicted under each plot. **B**. Example traces of each type of neuron from the modeled networks presented in *A*. **C**. Summation of the number of action potentials fired (1s/bin) of phasic and delayed firing excitatory neurons during the stimulation time shown in *A*, in simulated normal (*control, green*) vs. injury conditions (*injury-like, orange*). **D**. Bar graphs depict AUC from the plots shown in *C*. Note the increase in activity during simulated noxious stimulation compared to baseline, and substantial increase in response in the simulated injury model.

The analysis of the firing of the individual neurons divided into subcategories according to their network effect (excitatory/inhibitory) and their firing pattern show that both disinhibition and simulated injury-models lead to increased firing of the tonic inhibitory neurons along with increased firing in excitatory phasic and delayed neurons (**Figure 7A, B**). We then compared the firing of the excitatory phasic and delayed neurons in simulated naïve and injury conditions. Our model predicts that in naïve conditions, the excitatory phasic but not delayed neurons increase their firing slightly (**Figure 7C, D**, *green*), while in the simulated injury conditions, both excitatory phasic and delayed neurons substantially increase their firing (**Figure 7C, D**, *orange*). Considering that the projection neurons are excitatory with either phasic or delayed firing pattern [19,34,56] the substantial increase in firing of these excitatory neuronal types in response to the same noxious heat stimulus suggest that inflammation-induced amplified input to the SDH produces increased output from the SDH network.

### The synergistic effect of the increased peripheral input and SDH disinhibition

Our experimental and computational results show that a decrease in the SDH inhibition contributes to the increase in SDH activity (**Figures 5 and 6**). On the other hand, our model predicts that the increased input from the peripheral afferents conferred upon the SDH network is sufficient to affect the SDH activity without accompanying changes in the inhibitory transmission (**Figure 4**). In this case, increased afferent input also enhances the firing of tonic, inhibitory neurons (**Figure 7 A, B**, *right*), likely amplifying SDH inhibition. We, therefore, hypothesize that if both these mechanisms occur during injury, the combination of the effects of increased afferent input and injury-mediated SDH disinhibition, which opposes increased afferent input-mediated amplification of inhibition, might be synergistic in increasing the SDH activity. Application of the increased afferent input onto the SDH model with the 70% disinhibition led to a network saturation (*not shown*), suggesting a synergistic effect of the increased afferent input. Disinhibition to 20% led to a substantial increase in the network activity (**Figure 8A**) which was larger than the sum of the effects of the 20% disinhibition and increased afferent input (**Figure 8B**). These results suggest that both increased afferent input and disinhibition synergistically lead to an inflammation-induced increase in SDH network activity. To predict the extent of the inhibition needed to potentiate the effect of the increased afferent input on the SDH output, we examined the maximal firing of the excitatory (single, phasic and delayed firing) and inhibitory (tonic) neurons in “injury-like” conditions alone and compared how their firing changes when we add various degrees of disinhibition.

**Figure 8.**
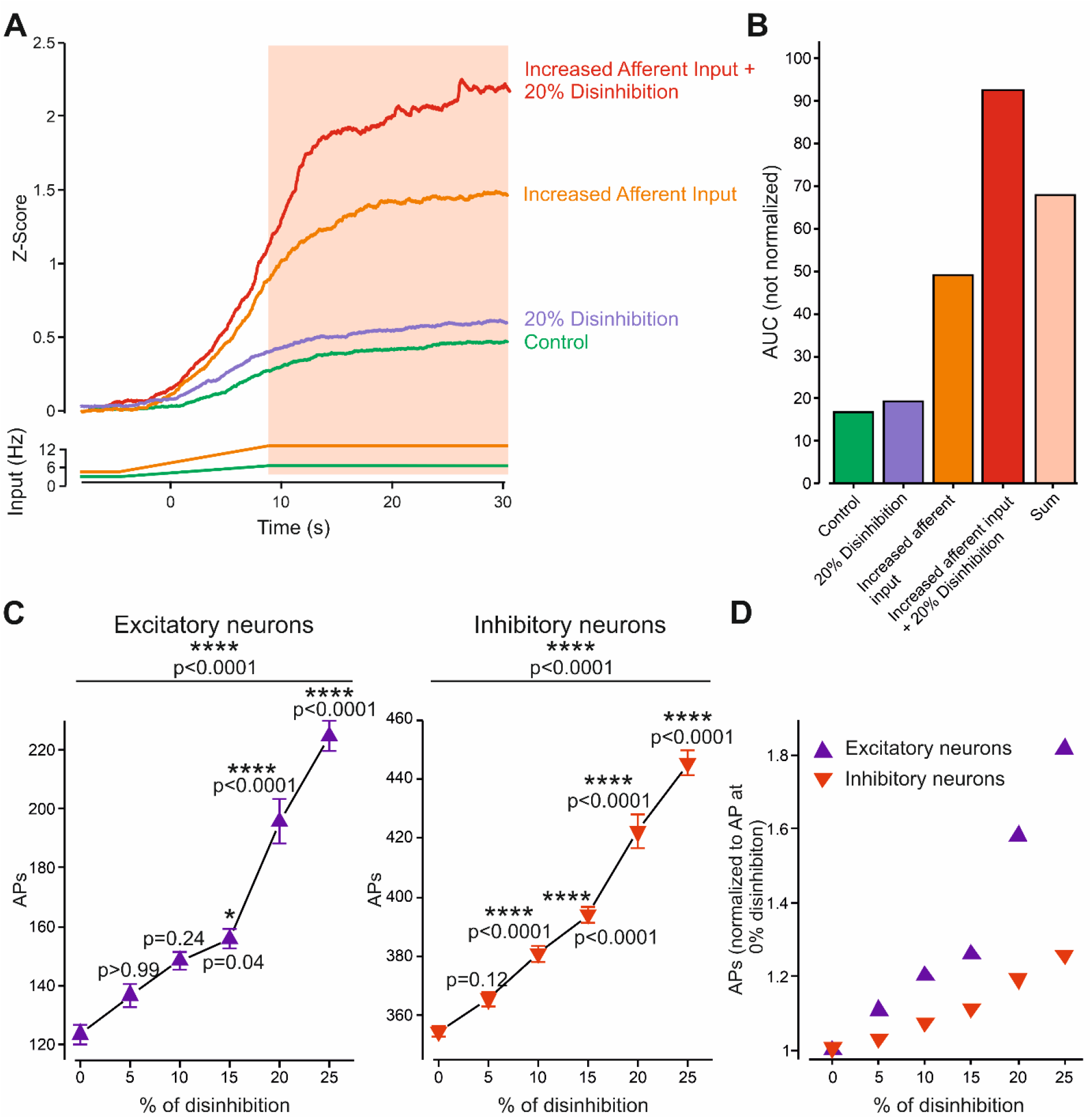
Disinhibition potentiates the effect of the increased afferent input on the SDH activity. **A**. Mean neural activity integration (1000 modeled neurons) in response to a simulated noxious heat stimulus applied in control (*green*), 20% disinhibition (*magenta*), increased afferent input (injury-like, *orange*), and increased afferent input applied to the network with 20% disinhibition (*red*) modeled conditions. The “raw” (not Z-scored) changes are presented, but all the traces are adjusted to a common baseline for easier comparison. **B**. Bar graphs depicting AUC of responses shown in *A*. The *pink* bar depicts the mathematical sum of the AUC values of 20% disinhibition and increased afferent input conditions. The AUCs are calculated from the “raw” data. **C**. Changes in AP firing of modeled excitatory (*left*) and inhibitory (*right*) neurons in response to constant increased afferent input (“injury-like” conditions) applied on the modeled SDH network with various degrees of disinhibition (in %). The 0% disinhibition shows AP firing in response to increased afferent input only. For the excitatory neurons, the AP firing of single, phasic and delayed firing neurons are summated. Means ± SD of the firing in 1 s bins along the time of stimulation (10 bins, pink shadow shown in *A*). RM two-way ANOVA with post-hoc Bonferroni, comparing different degrees of disinhibition to the AP firing in response to increased afferent input only (0% disinhibition). **D**. The increase in firing of excitatory (*blue triangles*) and inhibitory neurons (*red inverted triangles*) normalized to AP firing at 0% disinhibition.

We show that 10% of disinhibition is sufficient to increase the firing of the inhibitory neurons and 15% is sufficient to significantly increase the firing of the excitatory neurons compared to the increased afferent input state alone (0% disinhibition, **Figure 8C**). These data suggest that even a small decrease in inhibition is sufficient to potentiate the amplifying effect of increased afferent input on the SDH output. Interestingly, the relative increase in firing of the excitatory neurons following disinhibition was higher than the increase in firing of the inhibitory neurons (**Figure 8D**), suggesting that disinhibition primarily affects the firing of excitatory neurons. Altogether our results show that increased afferent input in synergism with the decrease in inhibition may underlie the injury-mediated increase in SDH activity and its output. The increased SDH output in response to the same noxious heat stimulus changes noxious heat stimulus perception, which is relayed to higher brain centers generating pain hyperalgesia.

## Discussion

A fundamental question in understanding pathological pain is what are the changes occurring within CNS neuronal networks that lead to an increased response to noxious heat stimuli and thereby amplifying pain [31]. Activation of specific receptors and ion channels in peripheral nerve endings by chemical, mechanical or thermal stimuli triggers the AP firing, generating the output of nociceptive neurons towards the CNS [3,17,27,66]. These input-output parameters of peripheral nociceptive neurons are altered under pathological conditions such as inflammation, which leads to changes in nociceptive intrinsic excitable properties, thus altering the transmission code by changing the rate and timing of the firing [3,7,10,70]. Here we utilized *in vivo* 2PLSM imaging of a large population of neurons to examine how CNS SDH circuitry changes its responses to abnormal peripheral inputs in inflammatory conditions. Our approach demonstrated how the alterations in the activity patterns of multiple neurons during the application of a noxious stimulus contribute to the overall activity amplification following injury. However, it did not consider the possible changes in the activity of the peripheral neurons following the end of the stimulation [34]. The alterations of these after-discharges may also contribute to the changes in the SDH activity. Our *in vivo* experiments also disregarded the inflammation-induced changes in the spontaneous activity of the peripheral afferents, which may enhance SDH activity [68]. Indeed, our computational model, which accounts for the changes in spontaneous activity, demonstrates that the increase in spontaneous activity substantially contributes to the overall inflammation-mediated changes in the SDH activity. We used the computational model we build to predict the possible mechanisms of the injury-mediated changes in the SDH activity. While our conclusions were robust using several combinations of parameters, we do not claim that the model is realistic in an absolute sense. For example, we used a current-based synapse that may not fully account for the voltage-dependent changes in the synaptic currents. However, our model was able to mimic the experimental results and recapitulate the same patterns of activity of the SDH neurons: activated, suppressed, and non-responsive.

It is noteworthy that we have performed our experiments in the young adult animals of 4-6 weeks at which the somatosensory system is not yet fully developed, however, it allowed easier tissue removal and, importantly, diminished attenuation of fluorescent signal due to relative tissue transparency at this age.

To detect the changes in the activity of SDH neurons, we loaded them with OGB. We chose OGB over more specific genetically encoded calcium indicators since we cannot find a proper transgenic mouse line that is pan-neuronally express GCaMP in SDH. The intra-spinal injections of the AAV expressing GCaMP are also not ideal as it leads to tissue inflammation, which might affect the properties of the spinal cord neurons and peripheral afferent axons [54, 55]. The disadvantage of using OGB is that it will measure the activity of neurons and astrocytes as well, implying that in our experiments, part of the recorded cells are astrocytes. However, since in our hands, only ~10% of all recorded cells were identified as astrocytes. This number may even be overestimated since SR101 has been shown to label SDH neurons [26]. Moreover, we show that SR101 labeled cells did not change their activity following injury. Considering all of the above, we have decided that the effect of the astrocytes on the overall activity is minimal, and therefore we linked the changes in the SDH activity to the alterations in the neuronal function.

Our in vivo results show that in normal conditions, the application of the noxious thermal stimuli leads to the increase in activity of only a fraction of neurons. About 30% of the recorded neurons remain silent following the application of noxious heat stimuli. It is feasible that SDH neurons that do not respond to the application of noxious heat stimuli are silenced by noxious-stimuli-triggered inhibition. This inhibition could result from activation of tonically active local inhibitory neurons and also from descending inhibition, triggered by noxious heat stimuli [45,62]. Our model containing only local inhibitory connections predicts that activation of local inhibitory SDH neurons by the noxious-like input in normal conditions is sufficient by itself to suppress and silence the activity of about 55% of modeled SDH neurons.

We show that this inhibitory drive is reduced following injury, which results in the de-suppression of the suppressed neurons and activation of the silenced neurons, thus contributing to the injury-mediated overall increase in the activity of the SDH network. What could be the origin of this disinhibition? Disruption of the chloride equilibrium or changes in the neuronal excitability and their synaptic efficacy were proposed as a principal mechanism underlying nerve-injury-mediated tactile allodynia [1,13,60,71]. The majority of these processes, however, require activation of complex cascades, which occur over a time scale of many hours [12,24]. Our experiments demonstrate the disinhibition-like changes of the SDH circuit that occur within 1 hour following injury. Therefore, it could be that the acute hyperalgesia we observed is mediated via a different, much more rapid mechanism. It is plausible that right after the injury, hyperexcited peripheral axons, similarly to the injured axons in nerve injury models [12], release a variety of mediators towards the SDH neurons, affecting the excitability of inhibitory neurons. The acute inflammation-mediated disinhibition will affect the activity of both inhibitory and excitatory neurons, but it would primarily enhance the activity of the excitatory SDH neurons as they rely more strongly on the inhibition to counterbalance the strong excitatory inputs they receive [32]. Indeed, our model predicts that disinhibiting the SDH would increase the firing of excitatory neurons by about 180%, whereas the firing of inhibitory neurons increases only by 125% (**Figure 8D**). Our experimental results further support the role of the disinhibition in the injury-induced increase in SDH activity by demonstrating that blockade of GABA and glycine receptors mimics the inflammation-induced SDH network activity changes, emphasizing yet again the role of spinal cord inhibitory transmission in controlling the output of the SDH network [8,9,24,60].

Our results show that increase in afferent input is sufficient to increase the SDH activity even without the changes in the SDH inhibition. The increased afferent input would amplify the activity of both excitatory and inhibitory neurons. It is, therefore, plausible that inflammation-triggered SDH disinhibition would potentiate the effect of increased afferent input on the excitation by reducing the effect of increased afferent input on the inhibition. Indeed, our computational model predicts that the combination of both: the inflammation-induced decrease in inhibition and inflammation-induced increase in afferent input synergistically enhances SDH output, thus encoding inflammatory hyperalgesia.

We show that the increased afferent input and SDH disinhibition alters the activity patterns of activated, suppressed, and NR neurons. The comparison of the number of activated, suppressed, and NR neurons that changed their activity patterns revealed a decrease in the fraction of initially activated, suppressed, and NR neurons that became suppressed following injury or disinhibition. These neurons became either NR or, in the case of initially suppressed neurons - activated. All these changes may contribute to the increase in the overall activity of the SDH network following injury or disinhibition. Interestingly, we show that injury and disinhibition lead to a decrease in the number of initially activated neurons that become suppressed following second noxious stimuli, which suggests that the activated neurons should also increase their activity following injury or disinhibition. The lack of the activity changes in the activated neurons measured *in vivo* could be due to a “saturation” or ceiling of the calcium signal such that the first noxious stimuli lead to substantial activation of the activated cells and that differences in AP firing following injury or disinhibition cannot be distinguished in calcium imaging. Alternatively, the lack of change in the activity of the activated neurons could be explained by the normalization of the higher baseline resulted from the inflammation- or disinhibition-mediated increase in the spontaneous activity (*see also an explanation on page 17*). Indeed, taking into account the changes in the baseline activity, the computational model of the SDH shows substantial activation of the activated cells following injury or disinhibition. Further analysis showed that the increased afferent input leads to the increased activity of the projection neurons. These results are in agreement with the previous studies showing increased firing of the projection neurons following acute inflammation[58].

In summary, here we introduced an approach that allows studying the SDH network representation of activity changes underlying pathological pain. By combining *in vivo* imaging with a network model of inflammatory pain, our data provide insight into the analysis of the representation of inflammatory hyperalgesia by spinal cord neurons. We show that inflammation-mediated nociceptive hyperexcitability is encoded by the acute changes in excitation-inhibition balance within the SDH, leading to a decrease in inhibition of activity and an increase in the number of activated neurons. These acute changes in the activity of pain-related spinal cord circuitry, by responding with higher excitatory output to the same noxious heat stimuli, might underlie changes in the perception of the noxious heat stimuli, thus leading to inflammatory hyperalgesia.

## Acknowledgements

We would like to thank Dr. Xioake Chen from Department of Biology, Stanford University and Dr. Qian Wang from Chen’s lab for training us in performing 2PLSM from SDH. We would also like to thank R. Sharon for the assistance with image registration performance. Support is gratefully acknowledged from the Canadian Institute of Health Research (CIHR), the International Development Research Centre (IDRC), the Israel Science Foundation (ISF) and the Azrieli Foundation - grant agreement 2545/18; Israeli Science Foundation - grant agreement 1470/17; the Deutsch-Israelische Projectkooperation program of the Deutsche Forschungsgemeinschaft (DIP) grant agreement B.I. 1665/1-1ZI1172/12-1 and Sessile and Seymour Alpert Chair in Pain Research.

## Supplementary Figures and Legends

**Supplementary Figure 1.**
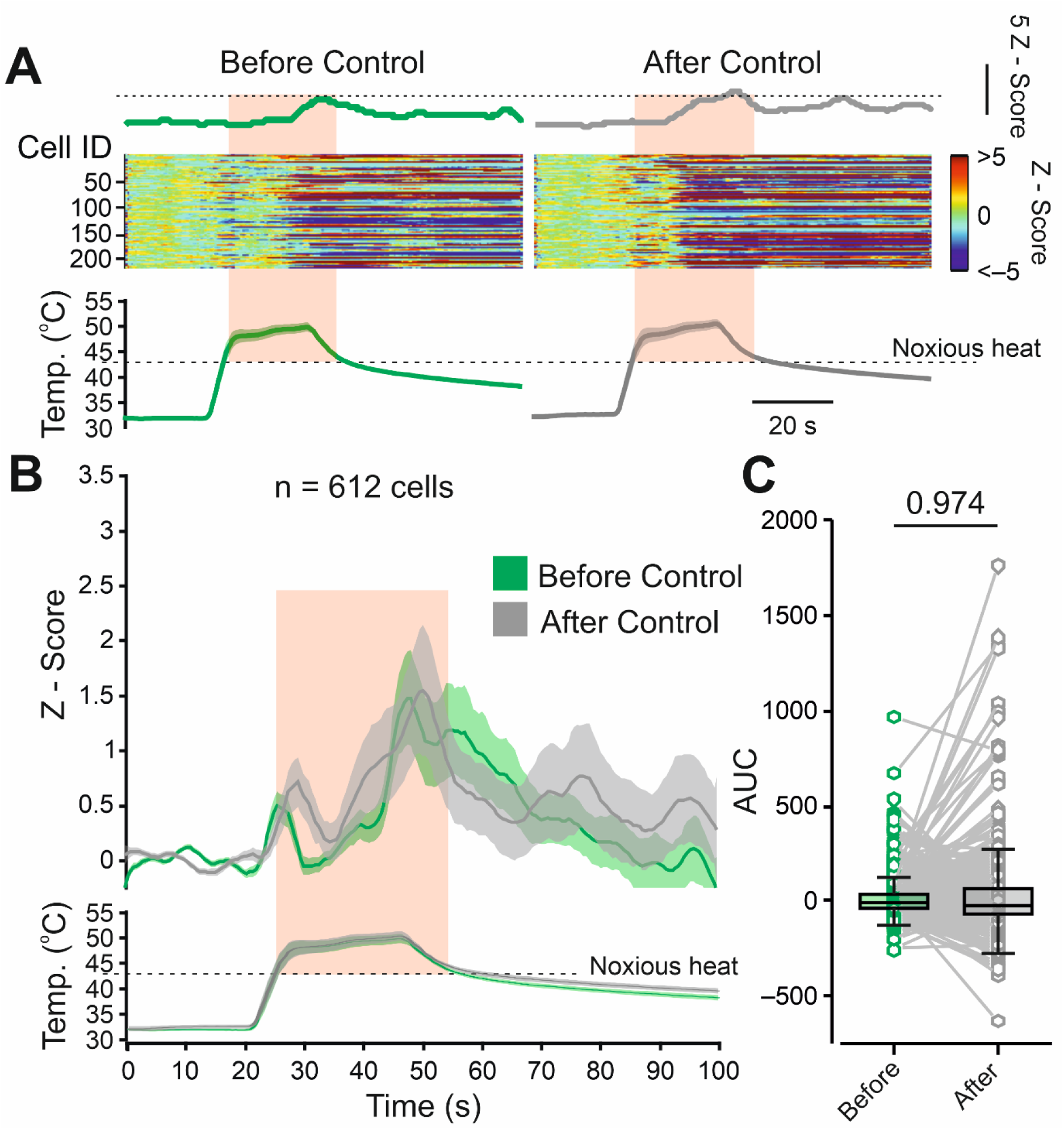
**A**. The representative activity heat maps for 209 cells in the same field of view (FOV) in control conditions, i.e., following application of first and second noxious heat stimuli, 1 hour apart without applying the hyperalgesia-inducing stimulus. Note that the response to the first noxious heat stimulus is similar to that of the second noxious heat stimulus. The averaged changes in activity are shown above the heat maps. The dotted line is aligned to the peak of the mean responses to the first and second noxious heat stimuli. The stimuli are shown below the heat maps, the noxious (above 42°C) component of the stimulus is outlined by the pink shadow. Representative of 4 mice. **B**. Changes in intracellular Ca^2+^ of all recorded cells in 4 mice in control condition, i.e., following application of first (*green*) and second noxious heat stimuli (*grey*), 1 h apart without applying the hyperalgesia-inducing stimulus. Note that the response to the first noxious heat stimulus is similar to that of the second noxious heat stimulus, suggesting the stability of the response. n=612 cells from 4 mice. **C**. Box plots and paired individual values of changes in the AUC of responses to the noxious heat in control conditions. The data points were collected and analyzed over the time of the noxious part of stimulus. Wilcoxon matched-pairs signed rank test, n=612 cells from 4 mice, Box plots depict the median and 25 and 75 percentiles, and the whiskers depict the outlier range.

**Supplementary Figure 2.**
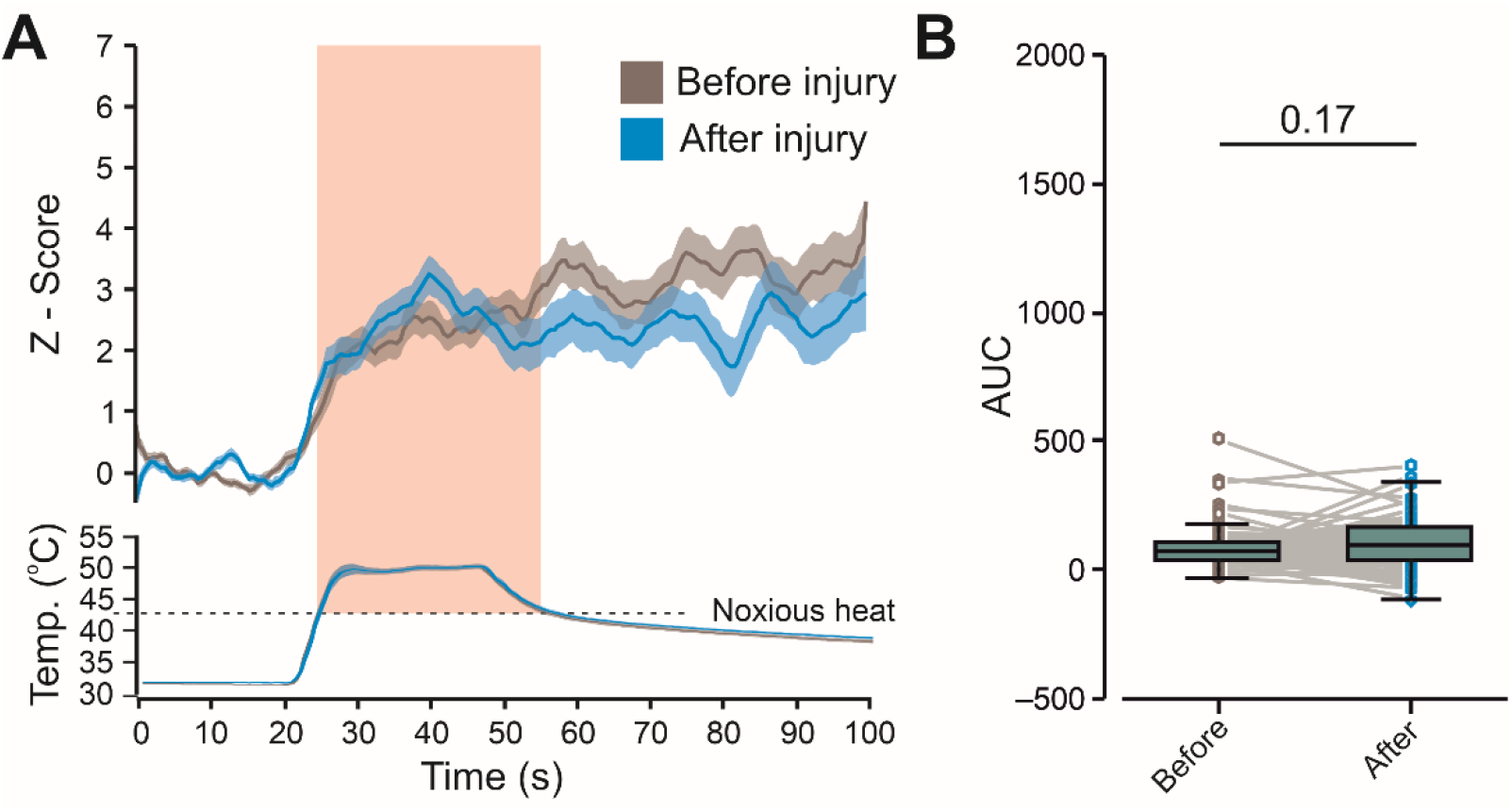
**A**. Changes in intracellular Ca^2+^ (Z-Score, mean ± SEM) in SR101-labeled astrocytes following application of the noxious heat stimulus (depicted below) before (*dark grey*) and 1 h after (*blue*) induction of the burn injury. The noxious (above 42°C) component of the stimulus is outlined by the pink shadow. n=77 astrocytes from 5 mice. **B**. Box plots and paired individual values of changes in AUC measured in astrocytes before (*dark grey*) and 1 hour after (*blue*) induction of the injury. The data points shown were collected and analyzed over the time of the noxious part of stimulus. Wilcoxon matched-pairs signed-rank test, n=77 astrocytes from 5 mice. Box plots depict the median and 25 and 75 percentiles, and the whiskers depict the outlier range.

**Supplementary Figure 3.**
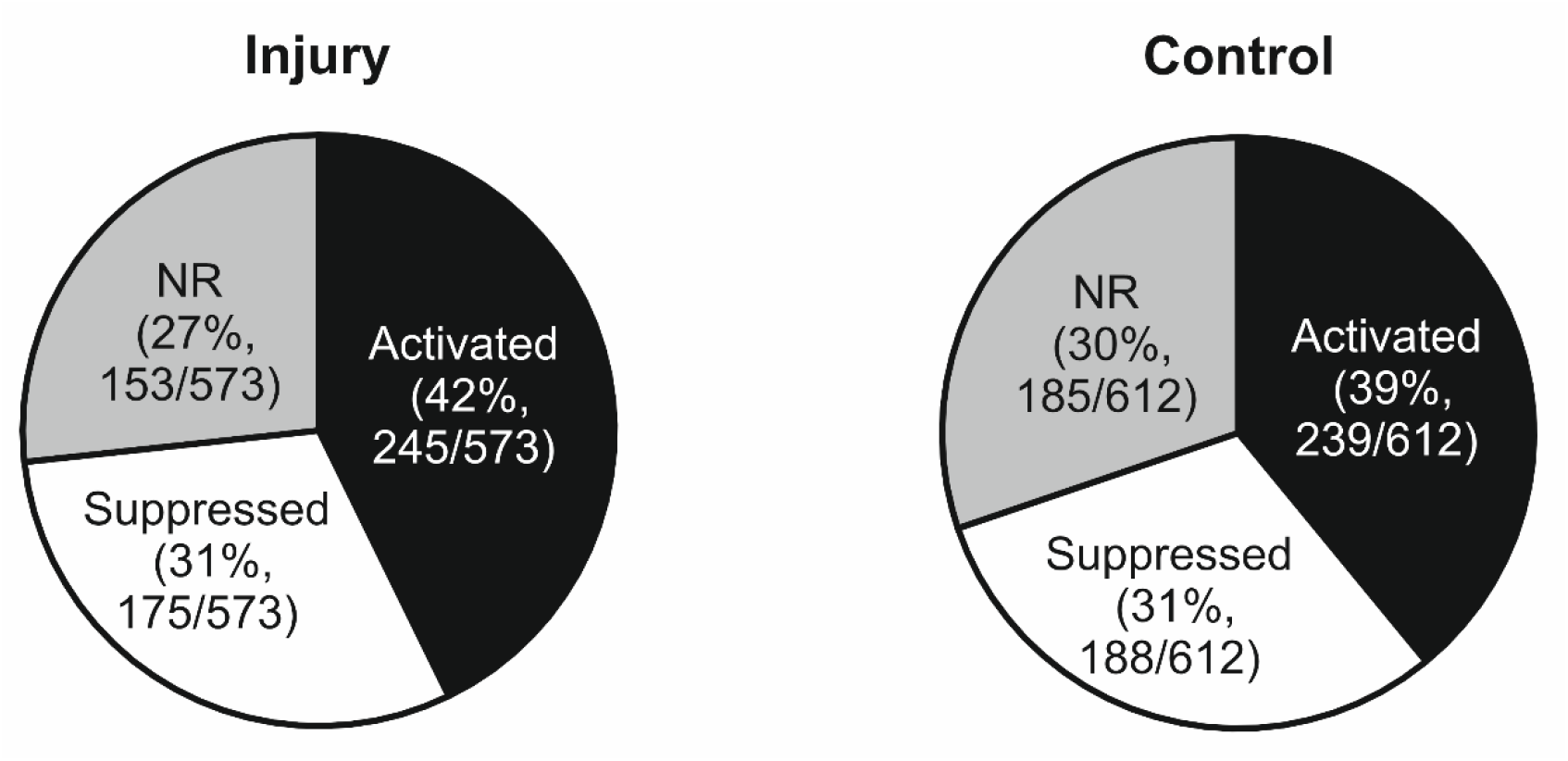
Pie charts of the fractions of NR, activated and suppressed cells in response to the application of the first noxious heat stimuli in the “injury” and “control” groups.

